# The hourglass organization of the *Caenorhabditis elegans* connectome

**DOI:** 10.1101/600999

**Authors:** Kaeser M. Sabrin, Yongbin Wei, Martijn van den Heuvel, Constantine Dovrolis

## Abstract

We approach the *C. elegans* connectome as an information processing network that receives input from about 90 sensory neurons, processes that information through a highly recurrent network of about 80 interneurons, and it produces a coordinated output from about 120 motor neurons that control the nematode’s muscles. We focus on the feedforward flow of information from sensory neurons to motor neurons, and apply a recently developed network analysis framework referred to as the “hourglass effect”. The analysis reveals that this feedforward flow traverses a small core (“hourglass waist”) that consists of 10-15 interneurons. These are mostly the same interneurons that were previously shown (using a different analytical approach) to constitute the “rich-club” of the *C. elegans* connectome. This result is robust to the methodology that separates the feedforward from the feedback flow of information. The set of core interneurons remains mostly the same when we consider only chemical synapses or the combination of chemical synapses and gap junctions. The hourglass organization of the connectome suggests that *C. elegans* has some similarities with encoder-decoder artificial neural networks in which the input is first compressed and integrated in a low-dimensional latent space that encodes the given data in a more efficient manner, followed by a decoding network through which intermediate-level sub-functions are combined in different ways to compute the correlated outputs of the network. The core neurons at the hourglass waist represent the *information bottleneck* of the system, balancing the representation accuracy and compactness (complexity) of the given sensory information.

**Author Summary:** The *C. elegans* nematode is the only species for which the complete wiring diagram (“connectome”) of its neural system has been mapped. The connectome provides architectural constraints that limit the scope of possible functions of a neural system. In this work, we identify one such architectural constraint: the *C. elegans* connectome includes a small set (10-15) of neurons that compress and integrate the information provided by the much larger set of sensory neurons. These intermediate-level neurons encode few sub-functions that are combined and re-used in different ways to activate the circuits of motor neurons, which drive all higher-level complex functions of the organism such as feeding or locomotion. We refer to this encoding-decoding structure as “hourglass architecture” and identify the core neurons at the “waist” of the hourglass. We also discuss the similarities between this property of the *C. elegans* connectome and artificial neural networks. The hourglass architecture opens a new way to think about, and experiment with, intermediate-level neurons between input and output neural circuits.

## Introduction

Natural, technological and information-processing complex systems are often hierarchically modular [1, 2, 3, 4]. A modular system consists of smaller sub-systems (modules) that, at least in principle, can function independently of whether or how they are connected to other modules: each module receives inputs from the environment or from other modules to perform a certain function [5, 6, 7]. Modular systems are often also hierarchical, meaning that simpler modules are embedded in, or reused by, modules of higher complexity [8, 9, 10, 11]. It has been shown that both modularity and hierarchy can emerge naturally as long as there is an underlying cost for the connections between different system units [12, 13].

In the technological world, modularity and hierarchy are often viewed as essential principles that provide benefits in terms of design effort (compared to “flat” or “monolithic” designs in which the entire system is a single module), development cost (design a module once, reuse it many times), and agility (upgrade, modify or replace modules without affecting the entire system) [14, 15, 16]. In the natural world, the benefits of modularity and hierarchy are often viewed in terms of evolvability (the ability to adapt and develop novel features can be accomplished with minor modifications in how existing modules are interconnected) [17, 18, 19] and robustness (the ability to maintain a certain function even when there are internal or external perturbations can be accomplished using existing modules in different ways) [20, 21, 22].

It has been observed across several disciplines that hierarchically modular systems are often structured in a way that resembles a *bow-tie or hourglass* (depending on whether that structure is viewed horizontally or vertically) [23, 24]. This structure enables the system to generate many outputs from many inputs through a relatively small number of intermediate modules, referred to as the “knot” of the bow-tie or the “waist” of the hourglass. The “hourglass effect” has been observed in systems of embryogenesis [25, 26], metabolism [27, 28], immunology [29, 30], signaling networks [31], vision and cognition [32, 33], deep neural networks [34], computer networking [35], manufacturing [36], as well as in the context of general core-periphery complex networks [37, 38]. The few intermediate modules in the hourglass waist are critical for the operation of the entire system, and so they are also more conserved during the evolution of the system compared to modules that are closer to inputs or outputs [39, 40, 35]. Note that the two terms, bow-tie and hourglass, have *not* been always interchangeable in the network science literature. In particular, the term bow-tie has been applied even to networks for which the knot includes a large fraction of the network’s nodes [41, 42].

In this paper, we apply the hourglass analysis framework of [23] on the *C. elegans* connectome [43]. The *C. elegans* connectome can be thought of as an information processing network that transforms stimuli received by the environment, through sensory neurons, into coordinated bodily activities (such as locomotion) controlled by motor neurons [43]. Between the sensory and motor neurons, there is a highly recurrent network of interneurons that gradually transforms the input information to output motor activity. An important challenge in applying the analysis framework of [23] on *C. elegans* is that the former assumes that the network from a given set of input nodes (sources) to a given set of output nodes (targets) is a Directed Acyclic Graph (DAG). On the contrary, the *C. elegans* connectome includes many nested feedback loops between all three types of neurons. For this reason, we extend the methods of [23] in networks that may include cycles as long as we are given a set of sources and a set of targets. The key idea is to identify the set of *feedforward paths* from each source towards targets, and to apply the hourglass analysis framework on the union of such paths, across all sources.

Our main result is that the *C. elegans* connectome exhibits the hourglass effect. This result is robust to the “routing methodology” that separates the feedforward from the feedback flow of information. Further, we observe the hourglass architecture when we consider just chemical synapses, or the combination of the latter with gap junctions. On the contrary, appropriately randomized networks do not exhibit the hourglass property. We also identify the neurons at the “waist” of the hourglass. Interestingly, they are mostly the same set of interneurons that were previously shown, using a different analytical methodology, to constitute the “rich-club” of the *C. elegans* connectome [44]. We explain that these two network architectures, hourglass and rich-club, are *not* equivalent – and in fact the hourglass property of the *C. elegans* connectome is maintained even if we rewire the connections between core neurons so that they do not form a rich-club. The fact that the core interneurons also form a rich-club suggests that *they form an information processing bottleneck that integrates the compressed information from different sensory modalities*, before driving any higher-level neural circuits.

We explain the benefits of the hourglass architecture, in the context of neural information processing systems, using an encoder-decoder model that resembles recent architectures in artificial neural networks [34, 45]. The encoding component compresses the redundant stimuli provided by the sensory neurons into a low-dimensional latent feature space (represented by the core neurons at the hourglass waist) that encodes the source information in a more efficient manner. Then, the decoding component of the network combines those latent features, which represent intermediate-level sub-functions, in different ways to drive each output through the motor neurons. The toy-example of Figure 1 illustrates this idea using a Boolean circuit with five binary sources and five output functions.

**Figure 1:**
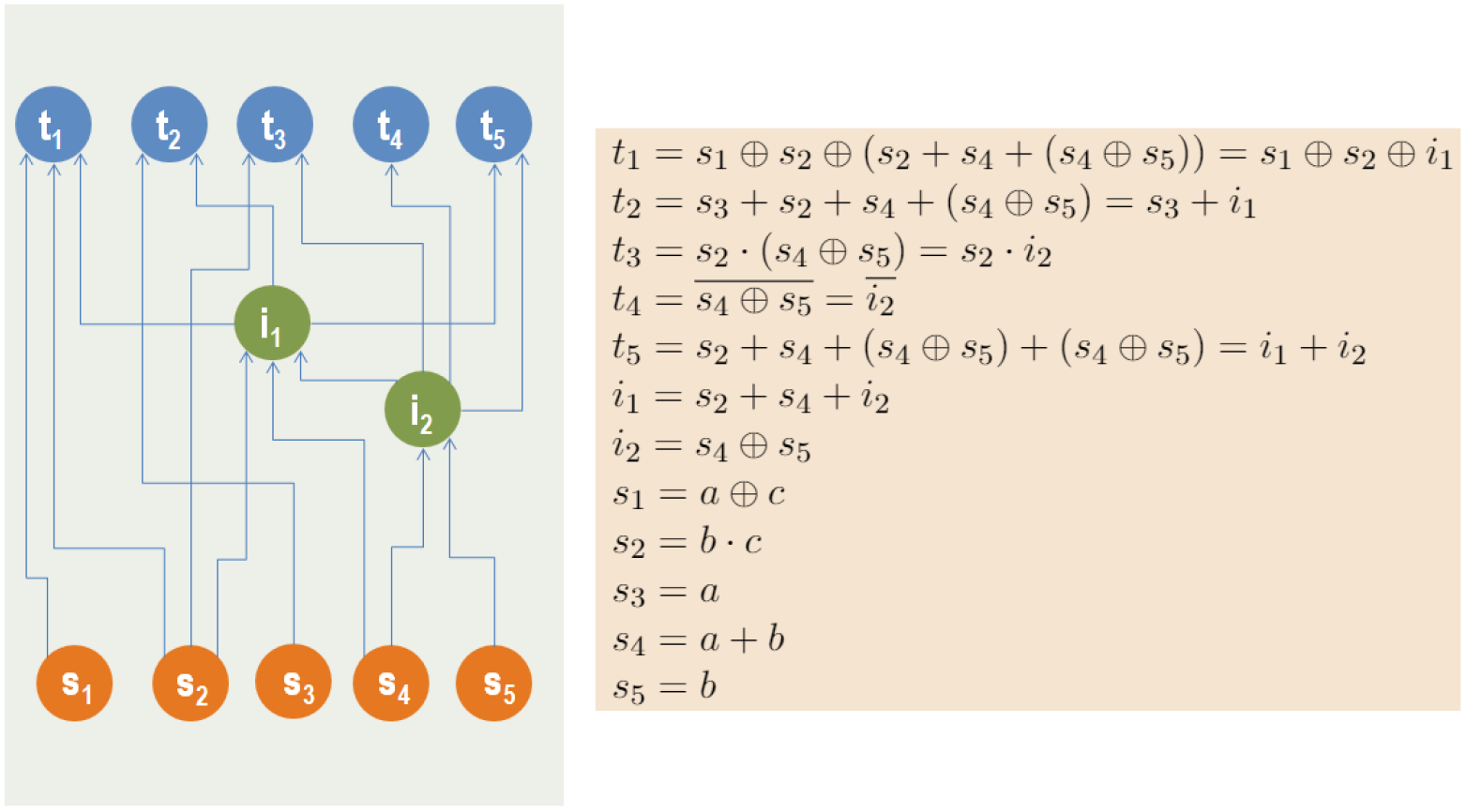
A hypothetical Boolean system with five sources and five targets. The sources are represented by orange nodes while the targets by blue nodes. Each target is a logic function of the sources. The sources are correlated, as shown by their logical expressions. A direct source-to-target computation would require 18 Boolean operations. Instead, we can compute the targets with only 9 operations if we first compute the two intermediate green nodes shown (3 operations) and then reuse those nodes to compute the targets (6 operations). This cost reduction is possible because there are correlations between the target functions. The two intermediate nodes, which represent the hourglass waist in this example, compress the information provided by the sources, computing sub-functions that are re-used at least twice in the targets. In this example the encoding part of the network is the set of connections between sources and intermediate nodes, while the decoding part is the set of connections between intermediate nodes and targets. In general, the encoder and decoder components can include additional nodes, creating a deeper hourglass architecture.

## Methods

### Connectome

The dataset we analyze describes the neural network of the hermaphrodite *C. elegans*, as reported in [43]. This connectome is a directed network between 279 neurons (the 282 non-pharyngeal neurons excluding VC6 and CANL/R, which are missing connectivity data). Neurons can be connected with two types of connections: chemical synapses and gap junctions (or, electrical synapses). The former are typically slower but strongest connections, and they transfer information only in one direction. The latter can be thought of as bi-directional connections.

The *synaptic network* (i.e., the network formed by only chemical synapses) consists of 2194 neural connections, created by 6393 chemical synapses. The *weight* of a connection is defined as the number of chemical synapses between the corresponding pair of neurons. The *in-strength* or *out-strength* of a neuron is defined as the sum of connection weights entering or leaving that neuron, respectively.

The *complete network* includes both chemical synapses and gap junctions. There are 514 pairs of neurons connected through gap junctions, creating the same number of bi-directional connections between those neurons. Unless mentioned otherwise, we analyze the synaptic network. In the Section “Including Gap Junctions: the Complete Network”, we extend the analysis to consider the complete network, asking whether there are any major differences when we also consider gap junctions.

The *C. elegans* neurons can be classified as sensory (S), inter (I) and motor (M) neurons, based on their structure and function [46]. Sensory neurons transfer information from the external environment to the central nervous system (CNS). Motor neurons transfer information from the CNS to effector organs (e.g. glands or muscles). Interneurons process information within the CNS. The *C. elegans* connectome has 88 sensory neurons, 87 interneurons and 119 motor neurons. Some of these neurons however have a dual role: ten behave as S and M, two as S and I, and three as M and I. In our analysis, we consider the S-M and S-I dual-role neurons as sensory, and the M-I neurons as motor. Consequently, the final network consists of 88 sensory neurons, 82 interneurons, and 109 motor neurons.

We can think of *C. elegans* as an information processing system in which the *feedforward flow* of information, from sensory to motor neurons, transfers sensory cues from the environment to the CNS, processes those signals to extract actionable information, which is then used to drive the behavior/motion of the organism. This feedforward flow however is regulated by multiple feedback loops that transfer information in the opposite direction, as well as lateral connections between neurons of the same type.

The connections that we refer to as *feedforward (FF)* are those from S to I, I to M, and S to M neurons. In the opposite direction (i.e., from I to S, M to I, and M to S neurons) the connections are referred to as *feedback (FB)*. Connections between neurons of the same type (i.e., S to S, I to I, and M to M neurons) are referred to as *lateral (LT)*. In the synaptic network, there are 901 FF connections, 998 LT connections, and 295 FB connections. Figure 2 shows the breakdown of these connection types in the synaptic and complete networks.

**Figure 2:**
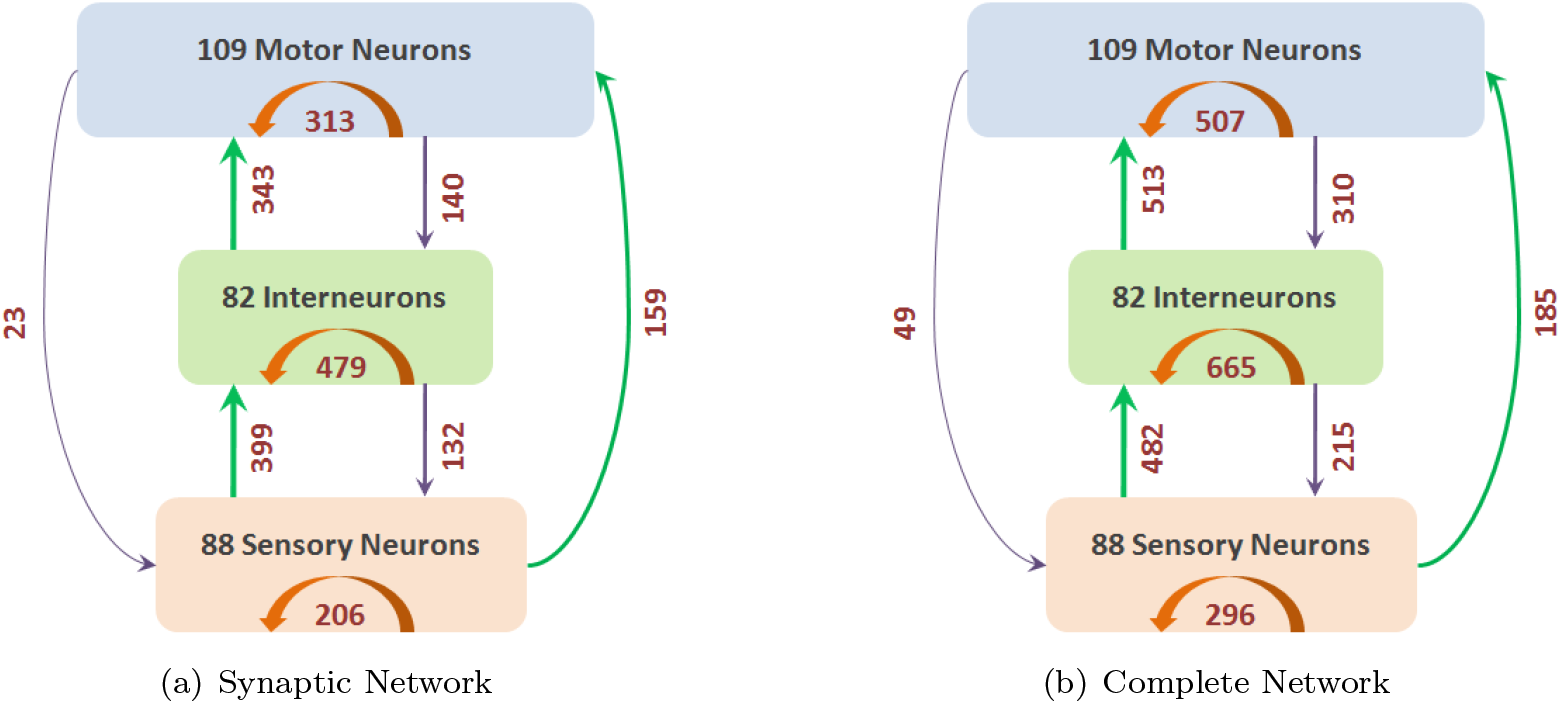
Neurons separated into three classes (S, I and M) based on structure and function. The number of connections between these three classes (and within each class) are shown with arrows (green for FF, orange for LT and purple for FB). (a) shows the synaptic network containing only chemical synapses, and (b) shows the complete network containing both chemical synapses and gap junctions.

The FB connection weights are often lower than FF and LT weights (see Figure 1(a)). Also, when considering neuron pairs that are reciprocally connected with both FF and FB connections, it is more likely that the FF connection is strongest than the corresponding FB connection (see Figure 1(b)). These observations suggest that the distinction between FF and FB connections has some neurophysiological significance.

If we focus on the top-5% stronger connections, relative to all chemical synapses, this set is dominated by feedforward S-to-I and I-to-M connections, as well as by lateral connections between I neurons and M neurons (see Table S1). None of the top-5% connections is of the feedback type. This observation suggests that feedback connections are weaker – one reason may be that they are involved mostly with the control of feedforward circuits, acting as modulators rather than drivers.

### Feedforward Paths from Sensory to Motor Neurons

The “routing problem” in a communication network refers to the selection of an efficient path, or multiple paths, from each source node to each target. In neural networks, there is no established “routing algorithm” that can accurately describe or model how information propagates from a sensory neuron to a motor neuron. Whether a neuron will fire or not depends on how many of its pre-synaptic neurons fire, the timing of those events, the physical size and location of the synapses in the dendritic tree, and several other factors. There are some first principles, however, that we can rely on to identify plausible routing schemes [47, 48]. These schemes should be viewed only as phenomenological models – we do not claim that neurons actually choose activation paths based on the following algorithms.

First, neurons cannot form routes based on information about the complete network or through coordination with all neurons (such as the routing algorithms used by the Internet or other technological systems). Instead, whether a neuron fires or not should be a function of only locally available information. So, we cannot expect that neural circuits use optimal routes that minimize the path length (“shortest path routing”) or other path-level cost functions [49].

Second, evolution has most likely selected routing schemes that result in efficient (even though not necessarily optimal) neural communication. Consequently, we can reject routing schemes that exploit all possible paths between two neurons as many of those paths would be inefficient.

Third, for robustness and resilience reasons, it is likely that multiple paths are used to transfer information from each sensory neuron to a motor neuron – schemes that only select a single path would be too fragile.

Fourth, given the low firing reliability of neurons, it is unlikely that a sensory neuron can communicate effectively with a motor neuron through multiple intermediate neurons. There should be a limit on the length of any plausible neural path [50].

Putting the previous four principles together, we are led to the following hypothesis: a sensory neuron S communicates with a motor neuron T through multiple paths that may be suboptimal but not much longer than the shortest path length from S to T.

Given this broad hypothesis, we identify several plausible routing schemes – and then examine whether our results are robust to the selection of a specific routing scheme.

To help choose reasonable parameter values for the various routing schemes we consider, we first examine the length and number of shortest paths from each sensory neuron S to each motor neuron M. Figure 2(a) shows the distribution of the length of these paths, measured in “hops” (i.e., connections between neurons). Almost all shortest paths from S to M neurons are between 2-4 hops. So, if the shortest connection from a sensory to a motor neuron is say 3 hops, the second and fourth principles suggest that we may also consider slightly longer paths, say 4 or 5 hops long.

Figure 2(b) shows the likelihood that an (S,M) pair is connected through *x* shortest paths. Note that only 4% of (S,M) pairs are not connected by any path, about 32% of (S,M) pairs are connected through only one shortest path, while the rest are connected with multiple shortest paths.

The various routing schemes we consider in the rest of the paper are (see Figure 3):

1. “*SP*”: As a reference point, *SP* refers to the selection of only shortest paths from a sensory neuron *s* to a motor neuron *t*.
2. “*SP*_4_”: The subset of *SP* including paths that are at most 4 hops.
3. “*SP*_5_”: The subset of *SP* including paths that are at most 5 hops.
4. “*SP* ^+1^”: The paths in *SP* together with all paths that are one hop longer than the shortest path from *s* to *t*.
5. “*SP* ^+2^”: The paths in *SP* together with all paths that are one or two hops longer than the shortest path from *s* to *t*.
6. 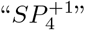: The subset of *SP* ^+1^ including paths that are at most 4 hops.
7. 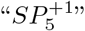: The subset of *SP* ^+1^ including paths that are at most 5 hops.
8. 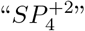: The subset of *SP* ^+2^ including paths that are at most 4 hops.
9. 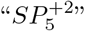: The subset of *SP* ^+2^ including paths that are at most 5 hops.
10. “*P*_4_”: All paths from *s* to *t* that are at most 4 hops long.
11. “*P*_5_”: All paths from *s* to *t* that are at most 5 hops long.

**Figure 3:**
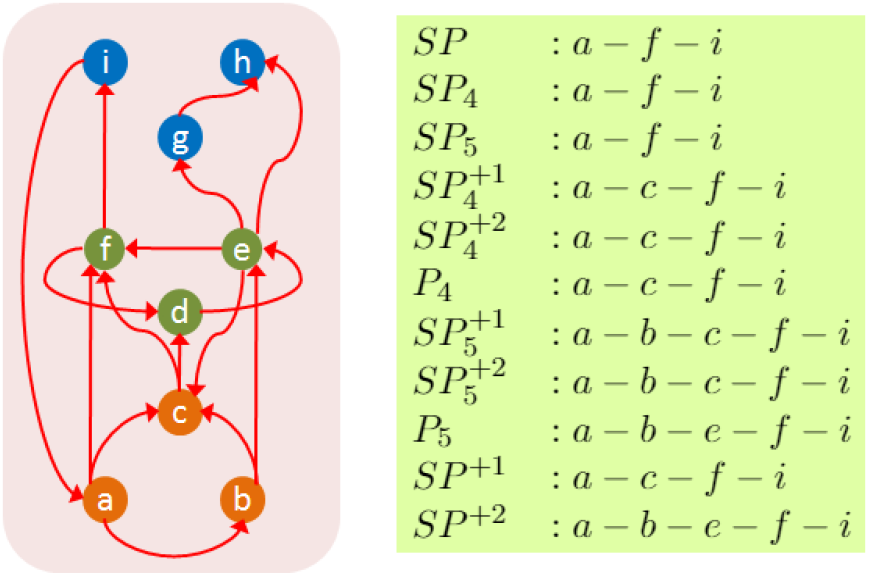
A representative path for each routing scheme when the source is *a* and the target is *i.* The orange nodes represent sensory neurons, the green nodes interneurons, and the blue nodes motor neurons.

The last two routing schemes (*P*_4_ and *P*_5_) are not variations of shortest path but they are based on the notion of diffusion-based routing. In the latter, information propagates from a source towards a sink selecting among all possible connections either randomly (e.g., random-walk based models) [51] or based on a threshold function (e.g., a neuron fires if at least a certain function of its pre-synaptic neurons fire) [52].

### Path Centrality Metric and *τ*-Core Selection

After utilizing one of the previous routing schemes to compute all paths from a sensory neuron to a motor neuron, we analyze these “source-target” paths based on the *hourglass framework*, developed in [23]. The objective of this analysis is to examine whether there is a small set of nodes through which almost all source-target paths go through. In other words, the hourglass analysis examines whether there is a small set of neurons that forms a bottleneck in the flow of information from sensory neurons towards motor neurons.

The path centrality *P* (*v*) of a node *v* is defined as the number of source-target paths that traverse *v*. This metric has been also referred to as the *stress* of a node [53]. Figure 4 illustrates the path centrality of each node in a small network – just for this example, the paths have been computed based on the shortest path (SP) routing algorithm. Any other routing scheme could have been used instead.

**Figure 4:**
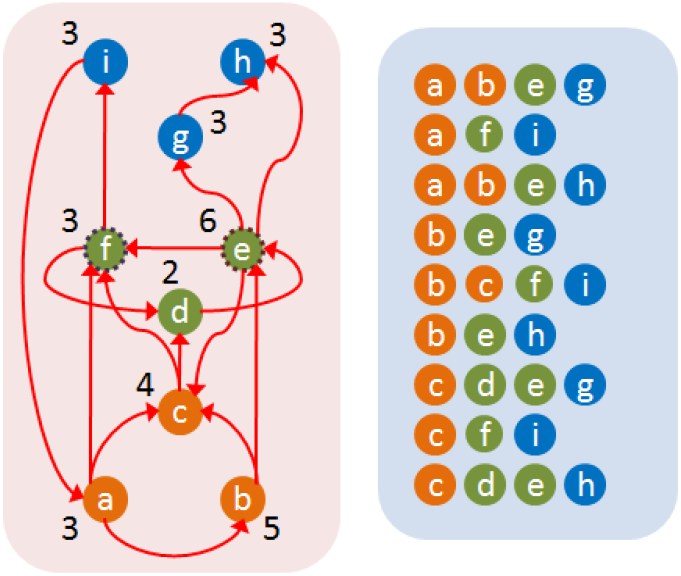
The path centrality of each node (shown at the left) based on the set of shortest paths *SP* (shown at the right). For *τ* =90%, a possible core is the two neurons {*e, f*}.

The path centrality metric is more general than betweenness or closeness centrality that are only applicable to shortest paths. Katz centrality does not distinguish between terminal and intermediate nodes and it penalizes longer paths. Metrics such as degree, strength, PageRank or eigenvector centrality are heavily dependent on the local connectivity of nodes rather than on the paths that traverse each node.

Given a set of source-target paths, the next step of the analysis is to compute the *τ-Core*, i.e., the smallest subset of nodes that can collectively cover a fraction *τ* of the given set of paths. The fraction *τ* is referred to as the *path coverage threshold* and it is meant to ignore a small fraction of paths that may be incorrect or invalid. Computing the *τ*-Core is an NP-Complete problem [23], and so we solve it with the following greedy heuristic (see [23] for an approximation bound):

- Initially, the core set is empty.
- In each iteration:

1. Compute the path centrality of all remaining nodes.
2. Include the node with maximum path centrality in the core set and remove all paths that traverse this node from the given set of paths.
- The algorithm terminates when we have covered at least a fraction *τ* of the given set of paths.

Figure 4 illustrates the core of a small network based on the shortest path routing mechanism, for *τ* =90%.

### Hourglass Score

Informally, the hourglass property of a network can be defined as having a small core, even when the path coverage threshold *τ* is close to one. To make the previous definition more precise, we can compare the core size *C*(*τ*) of the given network **G** with the core size of a derived network that maintains the same set source-target dependencies of **G** but that is not an hourglass by construction.

To do so, we create a *flat dependency network* **G**_**f**_ from **G** as follows:

1. **G**_**f**_ has the same set of source and target nodes as **G** but it does not have any intermediate nodes.
2. For every ST-path from a source *s* to a target *t* in **G**, we add a direct connection from *s* to *t* in **G**_**f**_. If there are *w* connections from *s* to *t* in **G**_**f**_, they can be replaced with a single connection of weight *w*.

Note that **G**_**f**_ preserves the source-target dependencies of **G**: each target in **G**_**f**_ is constructed based on the same set of “source ingredients” as in **G**. Additionally, the number of ST-paths in the original dependency network is equal to the number of paths in the weighted flat network (a connection of weight *w* counts as *w* paths). However, the paths in **G**_**f**_ are direct, without forming any intermediate modules that could be reused across different targets. So, by construction, the flat network **G**_**f**_ cannot have the hourglass property.

Suppose that *C*_*f*_ (*τ*) represents the core size of the flat network **G**_**f**_. The core of **G**_**f**_ can include a combination of sources and targets, and it cannot be larger than either the set of sources or targets. Additionally, the core of the flat network is larger or equal than the core of the original network (because the core of the flat network also covers at least a fraction *τ* of the ST-paths of the original network – but the core of the original network may be smaller because it can also include intermediate nodes together with sources or targets). So,

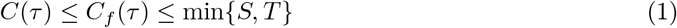

To quantify the extent at which **G** exhibits the hourglass effect, we define the *Hourglass Score*, or *H-score*, as follows:

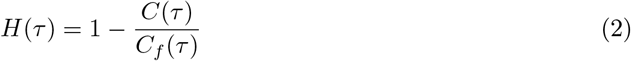

Clearly, 0 ≤ *H*(*τ*) < 1. The H-score of *G* is approximately one if the core size of the original network is negligible compared to the the core size of the corresponding flat network. Figure 5 illustrates the definition of this metric.

**Figure 5:**
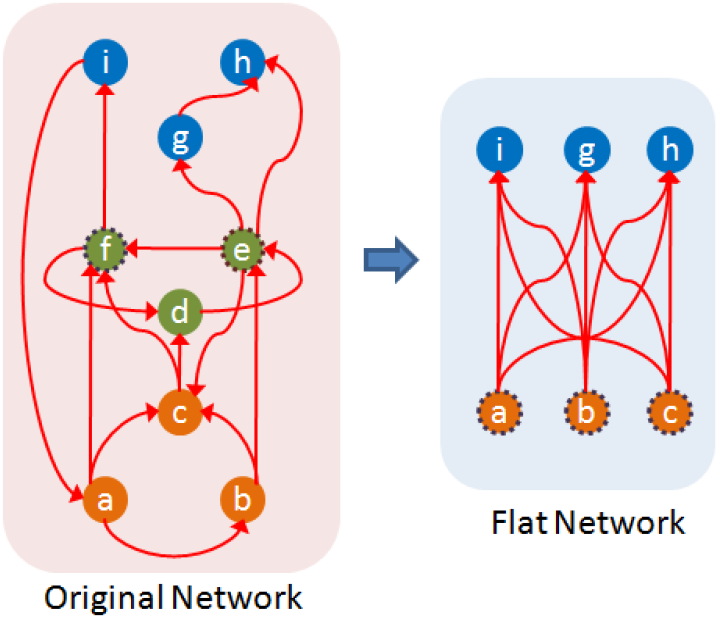
When the path coverage threshold is *τ* =90%, a core for the original network (left) is the set {*e, f*}. The weight of a connection in the flat network (right) represents the number of ST-paths between the corresponding source-target pair in the original network. All connections of this flat network have unit weight. The core of the flat network for the same *τ* consists of three nodes ({*a, b, c*} or ({*i, g, h*}). The H-score of the original network is 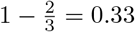.

An ideal hourglass-like network would have a single intermediate node that is traversed by every single ST-path (i.e., *C*(1)=1), and a large number of sources and targets none of which originates or terminates, respectively, a large fraction of ST-paths (i.e., a large value of *C*_*f*_(1)). The H-score of this network would be approximately equal to one.

### Randomization Method

We examine the statistical significance of the observed hourglass score in a given network *G* using an ensemble of randomized networks {*G*_*r*_}. The latter are constructed so that they preserve some key properties of *G*: the number of nodes and connections, the in-degree of each node, and the partial ordering between nodes (explained next). The randomization reassigns connections between pairs of nodes and changes the out-degree of nodes, as described below.

Suppose we are given *G* and a set of paths *P* from sources to targets. If there is a path in which node *v* appears after node *u and* there is no path in which *u* appears after *v*, we say that *u* is an ancestor of *v* and write *u* ∈ *A*(*v*). For a pair of nodes (*u, v*), we can have one of the following cases: (1) *u* is an ancestor on *v*, (2) *v* is an ancestor of *u*, (3) both *u* and *v* depend on each other, and (4) *u* and *v* do not depend on each other. We aim to preserve the partial ordering of nodes, as follows:

1. if *u* is *not* an ancestor of *v* in *G*, then it cannot be that *u* becomes an ancestor of *v* in a randomized network,
2. the set of ancestors of *v* in a randomized network is a subset of the set of ancestors *A*(*v*) in *G*.

The construction of randomization networks proceeds as follows: for each node *v* in the original network, we first remove all incoming connections. We then randomly pick in-degree(*v*) distinct nodes from *A*(*v*) and add connections from them to *v*. If in-degree(*v*) > |*A*(*v*)| then we add indegree(*v*) − |*A*(*v*)| additional connections (“multi-connections”) from randomly selected nodes in *A*(*v*) to *v*. The randomization mechanism is illustrated in Figure 6. It should be mentioned that there are several other randomization methods, preserving different network features [54]. None of them however preserve the partial ordering between nodes, which is an essential feature of a network in which a set of input-output dependency paths captures how information flows from sources to targets.

**Figure 6:**
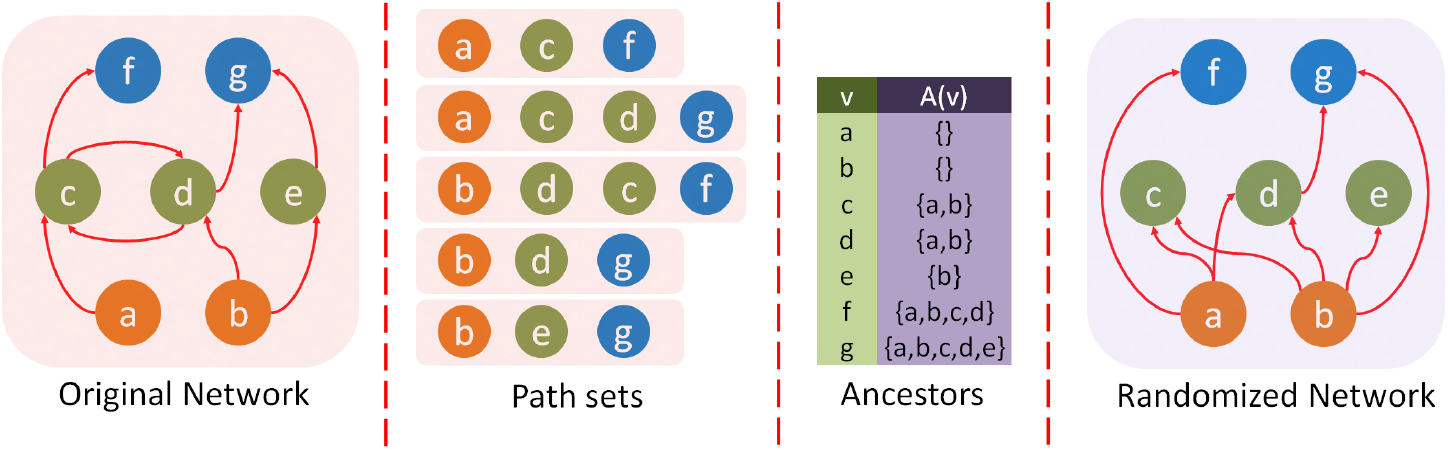
For the network and source-target paths given in the left two figures, we first compute the ancestors *A*(*v*) of each node *v* (shown in the third figure from l eft). In the randomized network (shown at the right), we preserve the in-degree of each node *v* and randomly select incoming connections from the set *A*(*v*).

### Location Metric

We also associate a *location* with each node to capture its relative position in the feedforward network between sources and targets. One way to place intermediate nodes between sources and targets is to consider the number of paths *P*_*S*_ (*v*) from sources (excluding *v* if it is a source itself) to *v* as a proxy for *v*’s *complexity* and the number of paths *P*_*T*_ (*v*) from *v* to targets (excluding *v* if it is a target itself) as a proxy for *v*’s *generality*. Nodes with zero in-degree (which cover most sources) have the lowest complexity value (equal to 0), while nodes with zero out-degree (which cover most targets) have the lowest generality value (equal to 0). The following equation defines a location metric based on *P*_*S*_ (*v*) and *P*_*T*_ (*v*),

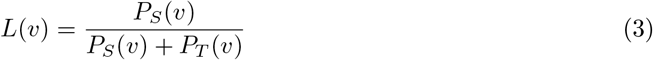

*L*(*v*) varies between 0 (for zero in-degree sources) and 1 (for zero out-degree targets). If there is a small number of paths from sources to a node *v* (low complexity) but a large number of paths from *v* to targets (high complexity), *v*’s role in the network is more similar to sources than targets, and so its location should be closer to 0 than 1. The opposite is true for nodes that have high complexity but low generality.

### Encoder-Decoder Architecture

Returning to the illustration of Figure-1, the number of paths from the set of sources *S* to a specific target *t* is denoted by *P*_*S*_ (*t*), and is equal to the number of source literals in the mathematical expression for *t*. If a Boolean expression involves *n* literals, it requires *n* − 1 Boolean operations. So, *P*_*S*_ (*t*) can be thought of as the *cost* for computing *t* from *S*.

More generally, even if a feedforward network does not represent Boolean expressions, we can think of the number of paths in *P*_*S*_ (*t*) as a cost metric for “computing” the target *t* from the set of sources *S*: the larger *P*_*S*_ (*t*) is, the more ways exist in which the information provided by the set of sources *S* affects the function of *t*.

Informally, an hourglass architecture is a network in which the information provided by a large set of sources *S* is first encoded (or compressed) into a small set *Z* of intermediate nodes at the “waist” of the hourglass. Then, the functions provided by the nodes in *Z* are decoded in computing the targets in *T*. Additionally, in an hourglass architecture there should be relatively few paths that bypass *Z*.

The question we focus on here is: *how does an hourglass architecture decrease the cost of computing a set of targets T from a set of sources S, and how large is that decrease in the case of C. elegans?*

Let *C*_*S*_ (*T*) be the cumulative cost for computing the set of targets *T* from the set of sources *S*:

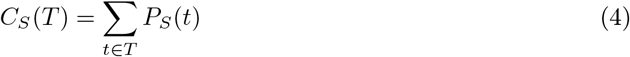

Given a set of intermediate nodes *Z*, we can produce the targets *T* in a two-step process: first, compute each node in *Z* from the sources *S*, and then compute each target in *T* from the set of intermediate nodes *Z*. There may be some source-to-target paths however that bypass the nodes in *Z* – we need to consider the cost of those “bypass-*Z*” paths as an extra term that depends on the selection of *Z*. So, the cost *C*_*S,Z*_ (*T*) of computing *T* from *S* given *Z* is:

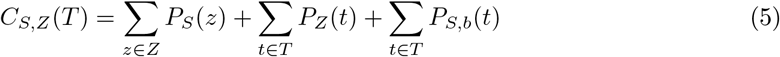

where the first summation term is the cost of computing *Z* from sources, the second is the cost of computing targets from *Z*, and the third is the cost of bypass-*Z* paths.

The *encoding-decoding gain* Φ_*Z*_, defined below, quantifies how significant is the cost reduction provided by such an encoder-decoder architecture,

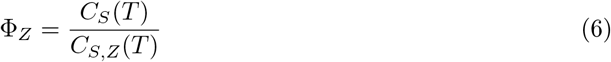

If Φ_*Z*_ ≤ 1, the intermediate nodes do not offer any cost reduction. On the other extreme, if there is a single intermediate node *z* that depends on all *n* sources and all *m* targets depend only on *z*, then Φ_*Z*_ gets its maximum value, *n m/*(*n* + *m*).

To illustrate, consider a three-layer network with *n* sources, *k* intermediate nodes and *m* targets, in which every intermediate node depends on every source, and every target depends only on every intermediate node. Suppose that the set *Z* consists of *k*′ < *k* of the intermediate nodes. It is easy to see that:

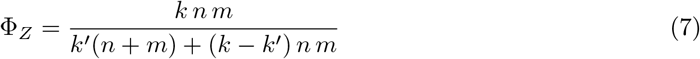

If *n* > 2 and *m* > 2, we have that *n* + *m* < *n m*, meaning that Φ_*Z*_ is maximized (equal to *n m/*(*n* + *m*)) when *Z* includes all *k* intermediate nodes (*k* = *k*′).

On the other hand, if the network includes *k*^+^ additional intermediate nodes that only connect to one source and one target, the maximum value of Φ_*Z*_ results when the set *Z* includes only the *k* densely connected nodes and leaves the *k*^+^ nodes in the bypass paths:

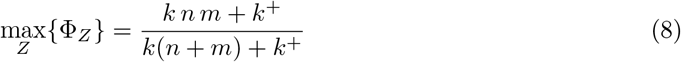

Returning to the network of Figure 1, the direct cost *C*_*S*_ (*T*) is ∑_*t*∈*T*_ *P*_*S*_ (*t*)=6+5+3+2+6=22. The cost of constructing the nodes in *Z* from sources is ∑_*z*∈*Z*_ *P*_*S*_ (*z*)= 4+2=6, the cost of constructing targets from *Z* nodes is ∑_*t*∈*T*_ *P*_*Z*_ (*t*)=1+1+1+1+2=6, while the cost of bypass-*Z* paths is ∑_*t*∈*T*_ *P*_*S,b*_(*t*)=2+1+1+0+0=4. So, the encoding-decoding gain is 22/16=1.375 while its maximum possible value is 25/10=2.5.

## Results

### Hourglass Analysis of Feedforward Paths

We defined earlier eleven different routing methods for computing paths from sensory to motor neurons in *C. elegans*. Table 1 shows some relevant properties for each of these path sets. The number of all possible pairs of sensory-motor neurons is about 9,500. About 90%-95% of these pairs are connected with any of the eleven path sets. Even with the smallest path set (*SP*_4_), there are typically multiple paths for every sensory-motor pair. The sensory-motor neural paths are typically short: the median is 3 hops meaning that there are typically two other neurons between a sensory and a motor neuron. The number of paths increases ten-fold when we allow one more hop than the shortest path (*SP* ^+1^), and about eight-fold more when we allow a second extra hop (*SP* ^+2^). Also, about 5%-10% of the connectome (after the removal of FB connections between the three different classes of neurons) are not traversed by any of these paths. This suggests that these connections are utilized only in feedback circuits between neurons of the same type (e.g., feedback between interneurons).

**Table 1:**
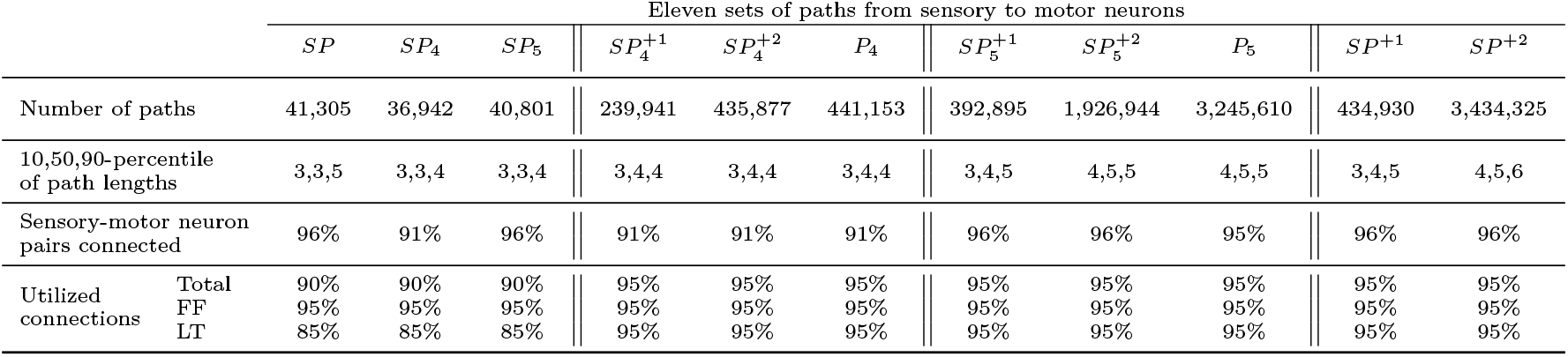
Properties of the eleven paths sets from sensory to motor neurons computed using the eleven routing methods we consider.

Given a set 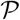 of feedforward paths from sensory to motor neurons, we now apply the hourglass analysis framework (see Section “Hourglass Score”). In particular, the goal is to compute the smallest set of neurons that can cover a percentage *τ* of all paths in 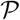. That set of neurons is referred to as *τ-Core*.

Figure 7(a) shows the cumulative path coverage as a function of the number of neurons in the *τ*-Core (increasing values of *τ* require a larger set of core neurons). All path sets have the same “sharp knee” property: almost all (80%-90%, depending on the routing method) of the feedforward paths traverse a small set of about 10 neurons. Routing methods that produce more paths (such as *SP* ^+2^) tend to have a smaller *τ*-Core than more constrained routing methods (such as *SP*_4_).

**Figure 7:**
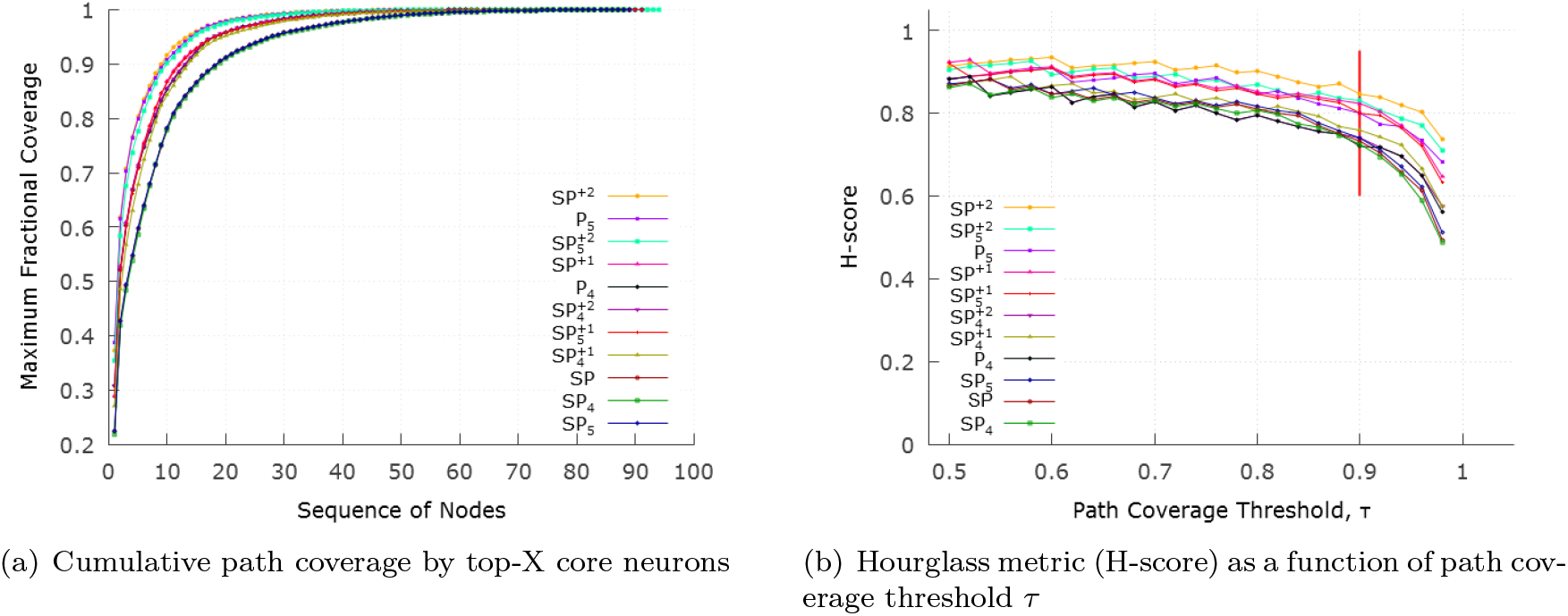
(a) Cumulative path coverage by the top-*X* core neurons for *X* = 1 … 100. Depending on path selection method, the core size varies between 10-20 neurons when the path coverage threshold *τ* is 90%. For a given *τ*, routing methods that produce more paths (such as *SP* ^+2^ or *P*_5_) result in a smaller core. (b) Effect of *τ* on hourglass metric (H-score) for each path set. The ordering of the legends is the same with the top-down ordering of the curves.

Figure 7(b) examines the effect of the path coverage threshold *τ* on the hourglass metric (H-score). For all path sets, the H-score is close to one (its theoretical maximum value) as long as *τ* < 90%. This suggests an hourglass-like architecture, independent of which routing scheme has produced the set of feedforward paths.

Table 2 shows the sequence of core neurons (for *τ* = 90%) for each path set. The first 10-11 of those neurons appear in almost every path set. The remaining neurons appear in more constrained path sets (such as *SP*) and they only cover a small fraction of additional paths (1%-3%).

**Table 2:** The identified core neurons when the path coverage threshold is *τ* = 90% for each path set. For each core neuron, we show the fraction of paths that the corresponding neuron contributes to the core. The neurons are ranked in decreasing order in terms of their contribution to the core (considering the *SP* set of paths), grouping bilateral neurons together. The last column shows the 11 “rich-club” neurons, as identified in [44].

| Core Neuron | $SP$ | $SP_4$ | $SP_5$ | $SP_4^{+1}$ | $SP_4^{+2}$ | $P_4$ | $SP_5^{+1}$ | $SP_5^{+2}$ | $P_5$ | $SP^{+1}$ | $SP^{+2}$ | Rich-club |
| --- | --- | --- | --- | --- | --- | --- | --- | --- | --- | --- | --- | --- |
| AVAL | 0.25 | 0.24 | 0.25 | 0.24 | 0.24 | 0.24 | 0.26 | 0.25 | 0.25 | 0.26 | 0.27 | ✓ |
| AVAR | 0.23 | 0.22 | 0.23 | 0.30 | 0.34 | 0.34 | 0.32 | 0.40 | 0.43 | 0.32 | 0.41 | ✓ |
| AVBL | 0.08 | 0.07 | 0.08 | 0.10 | 0.09 | 0.09 | 0.09 | 0.10 | 0.10 | 0.09 | 0.10 | ✓ |
| AVBR | 0.04 | 0.04 | 0.04 | 0.04 | 0.03 | 0.03 | 0.04 | 0.03 | 0.03 | 0.04 | 0.03 | ✓ |
| AVEL | 0.06 | 0.06 | 0.06 | 0.05 | 0.04 | 0.04 | 0.05 | 0.04 | 0.03 | 0.05 | 0.03 | ✓ |
| AVER | 0.06 | 0.06 | 0.06 | 0.06 | 0.05 | 0.05 | 0.06 | 0.05 | 0.04 | 0.05 | 0.04 | ✓ |
| DVA | 0.05 | 0.05 | 0.04 | 0.02 | 0.03 | 0.03 | 0.03 | 0.03 | 0.02 | 0.04 | 0.01 | ✓ |
| PVCL | 0.05 | 0.05 | 0.05 | 0.07 | 0.07 | 0.07 | 0.07 | 0.07 | 0.07 | 0.06 | 0.06 | ✓ |
| PVCR | 0.03 | 0.03 | 0.03 | 0.02 | 0.02 | 0.02 | 0.02 |  |  | 0.03 | 0.04 | ✓ |
| AVDR | 0.04 | 0.04 | 0.04 | 0.04 | 0.03 | 0.03 | 0.03 | 0.02 | 0.02 | 0.03 | 0.01 | ✓ |
| AVDL | 0.02 | 0.02 | 0.02 | 0.02 | 0.01 |  | 0.01 |  |  |  |  | ✓ |
| HSNR | 0.02 | 0.03 | 0.03 | 0.02 | 0.02 | 0.01 | 0.01 | 0.01 |  | 0.02 |  |  |
| HSNL | 0.01 |  | 0.01 |  |  |  |  |  |  |  |  |  |
| RIAL | 0.01 | 0.02 | 0.01 | 0.01 | 0.02 | 0.02 | 0.01 |  | 0.01 | 0.01 |  |  |
| RIAR | 0.01 | 0.02 | 0.01 |  |  | 0.01 |  |  |  |  |  |  |
| RIMR | 0.01 | 0.01 | 0.01 | 0.01 | 0.01 | 0.02 |  |  |  |  |  |  |
| RMGL | 0.01 | 0.01 | 0.01 |  |  |  |  |  |  |  |  |  |
| PVR | 0.01 | 0.01 | 0.01 |  |  |  |  |  |  |  |  |  |
| AIBR | 0.01 | 0.01 | 0.01 |  |  |  |  |  |  |  |  |  |
| AIZL |  | 0.01 |  |  |  |  |  |  |  |  |  |  |
| H-score | 0.79 | 0.79 | 0.79 | 0.82 | 0.84 | 0.74 | 0.85 | 0.86 | 0.81 | 0.84 | 0.87 |  |

If we focus on those first 10-11 core neurons, we observe that, first, they are included in the 90%-core of all path sets we consider (with few exceptions).

To simplify the presentation of the results, in the rest of this paper we will focus on the “*SP* ^+2^” path set. This path set results in the largest number of paths and a core of 10 neurons when *τ* =90%.

That set of core neurons includes bilateral pairs of interneurons (namely: AVA, AVB, PVC, AVE, and AVD) – the DVA stretch sensitive core neuron does not appear bilaterally. Seven of the core neurons are located in the head region (AVAR/L, AVBR/L, AVER/L, AVDR) and three are in the tail region (PVCR/L, DVA). The original ten core neurons contain nine command interneurons that play a pivotal role in forward and backward locomotion [44]. The other non-command interneuron of the core, DVA, is a proprioceptive interneuron modulating the locomotion circuit [44].

If we want to extend the set of core neurons slightly by covering *τ* =95% of all paths instead of 90%, we need to add four more neurons into the core (HSNR, AVDL, RIAL, RIMR).

### Comparison with Rich-Club Effect

The existence of a set of densely interconnected nodes in the *C. elegans* connectome, termed as “rich-club”, has been previously established by Towlson et al. [44]. A rich-club is a subgraph of high-degree nodes that are much more densely interconnected with each other than what would be expected based only on their degrees [55]. In other words, the rich-club concept is based on the analysis of local connectivity in a network – rather than the analysis of (shortest or other) network paths. Further, the rich-club analysis does not consider whether some nodes act as inputs (sensory neurons) or outputs (motor neurons) in the network. The hourglass analysis, on the other hand, analyzes the set of feedforward paths from inputs to outputs. So, these two methods are significantly different.

Are these two network properties, rich-club and hourglass effect, equivalent? We can see that this is not the case through simple counter-examples (see Figure 8(a)).

**Figure 8:**
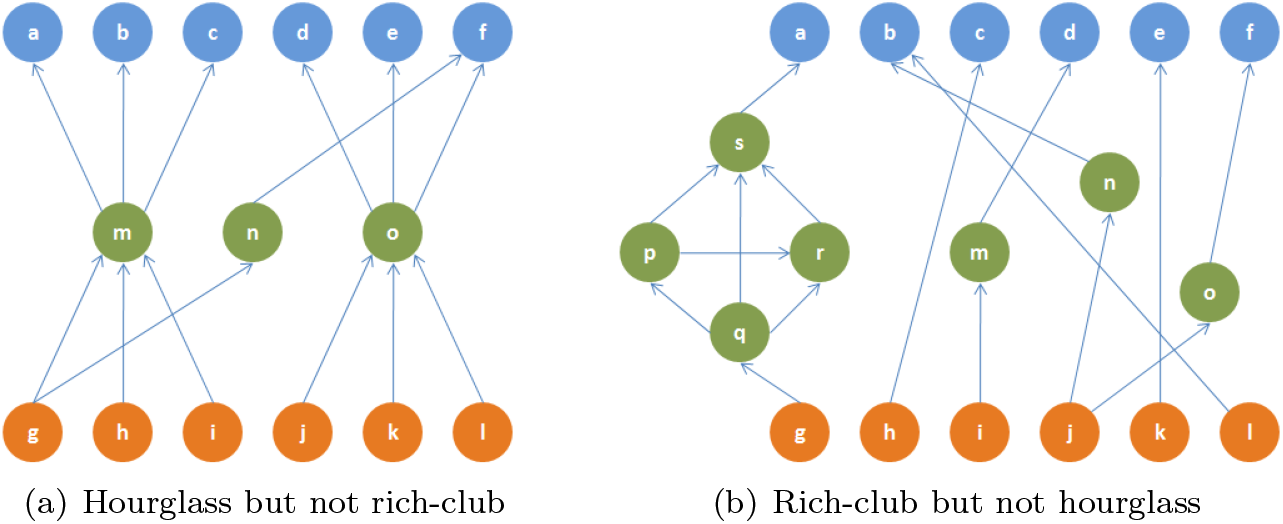
(a) A toy network in which two nodes (*m* and *o*) cover more than 90% of all source-target paths (*H*-score=0.67). This network does not contain a rich-club. (b) A toy network that is not an hourglass (*H*-score=0) but it has a rich-club (nodes *p, q, r, s* – the rich-club coefficient is 2.60 [56]).

An important observation, however, is that the core neurons that we identify through the hourglass analysis highly overlap with the rich-club neurons of [44]. The first ten core neurons identified by *all* routing methods we consider also appear in the eleven rich-club neurons reported in [44]. The AVDL interneuron is the 11th rich-club member but it appears in the hourglass core only in half of the routing methods we consider (for *τ* =90%). The fact that two very different methods highlight almost the same set of interneurons as the most important in the system adds confidence in the results of both studies.

The fact that a small set of interneurons act as *both* the hourglass core and rich-club, even though these two network properties are qualitatively different, raises an interesting hypothesis about the functional role of these interneurons: In the hourglass network of Figure 8(a), the core nodes *m, n, o* are *not* connected with each other – such an architecture can compress different input information streams but without integrating them. On the contrary, the core interneurons of *C. elegans* are densely interconnected and so *they form an information processing bottleneck that integrates the compressed information from different sensory modalities*, before driving any higher-level neural circuits.

### Comparison with Randomized Networks

Is the hourglass effect a genuine property of the *C. elegans* connectome or would it also be present in similar but randomly connected networks? We generate 1000 random networks using the algorithm described in Section “Randomization Method”. The randomization process preserves the in-degree of each neuron and the hierarchical ordering between neurons (i.e., if neuron *v* depends on neuron *u* but *u* does not depend on *v* in the original connectome, it cannot be that *u* depends on *v* in a randomized network). Figure 9 shows the H-score distribution of the randomized networks. The H-score of the random networks is significantly less than the corresponding original network (*p* < 10^−3^), suggesting that the hourglass effect we observe in the *C. elegans* connectome is not a statistical artifact.

**Figure 9:**
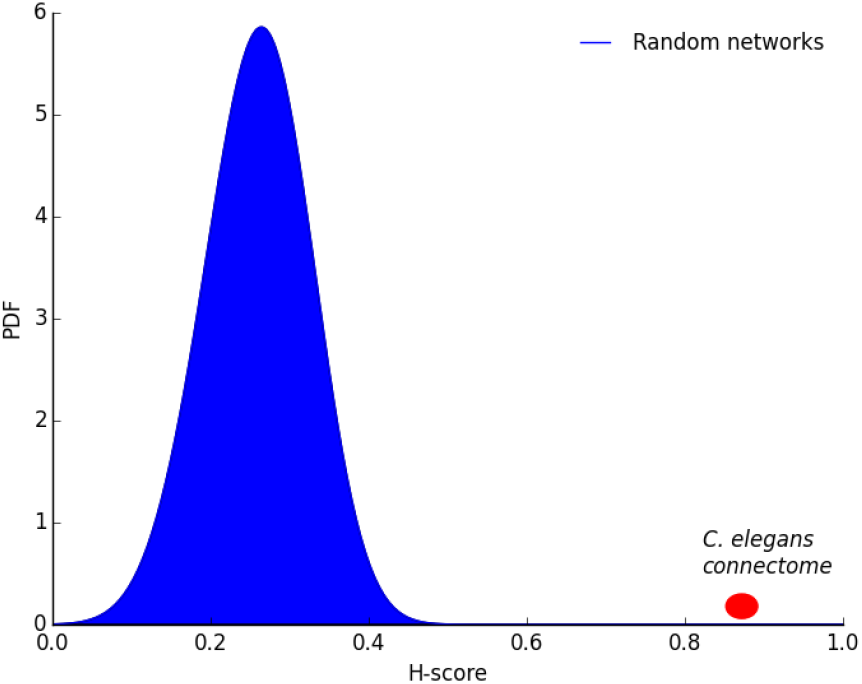
Distribution of H-score for randomized networks in which we preserve the in-degree of each neuron and the hierarchical ordering between neurons. The probability of observing the H-score value of the original network in randomized networks is less than 10^−3^.

Is the hourglass effect a consequence of the dense connectivity between core neurons? The latter is the defining characteristic of rich-club neurons. Would we still observe the hourglass effect if the core neurons were not so densely interconnected with each other, forming a rich-club?

To answer this question, we perform a second randomization experiment in which every connection between two core neurons *X* and *Y* is rewired so that *X* connects instead to a randomly chosen neuron *Z* that is *not* in the set of core neurons. We experimented with two variations of this method: one in which *Z* is an interneuron and another in which *Z* can be any neuron, including sensory and motor neurons.

Both approaches *fail* to destroy the hourglass property. As shown in Figure 10, the *H*-score distribution of the randomized networks (100 instances) includes the *H*-score of the original network (0.87). This means that the hourglass property is *not* due to the dense connectivity between core neurons. When we remove the connections between core neurons, we reduce the number of core nodes that a typical sensory-to-motor path traverses – but it is still the case that almost all such paths traverse at least one core node, and this is what creates the hourglass property.

**Figure 10:**
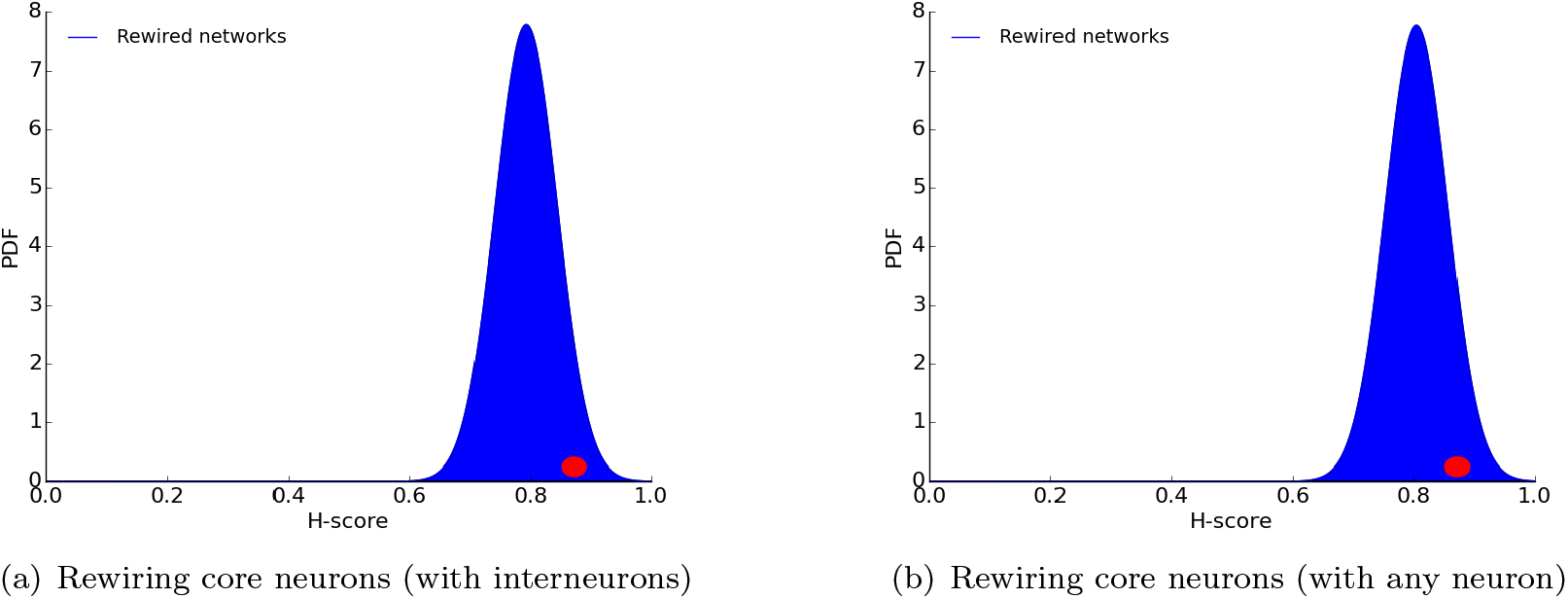
H-score distribution of randomized networks in which every connection *X*-*Y* between two *core* neurons *X* and *Y* is rewired. In (a), *Y* is replaced with a randomly chosen *interneuron Z* that is not in the core. In (b), *Y* is replaced with a randomly chosen neuron *Z* (including sensory and motor neurons) that is not in the core. The red dot shows the *H*-score of the original connectome.

### Hourglass Organization based on Location Metric

The location metric associates each neuron *v* with a value between 0 and 1, depending on the number of paths from sensory neurons to *v* and from *v* to motor neurons.

Figure 11 shows the location of each neuron in a vertical orientation. Most sensory neurons have zero incoming connections (and thus no incoming paths), and so their location is 0. Similarly most motor neurons have zero outgoing connections and so their location is 1. The location of the ten core neurons is shown with dotted rectangles – they are concentrated close to the middle of the location range, meaning that their number of paths from sensory neurons is roughly the same with their number of paths to motor neurons. This visualization has been produced with the *SP* ^+2^ path set but other path sets give similar results.

**Figure 11:**
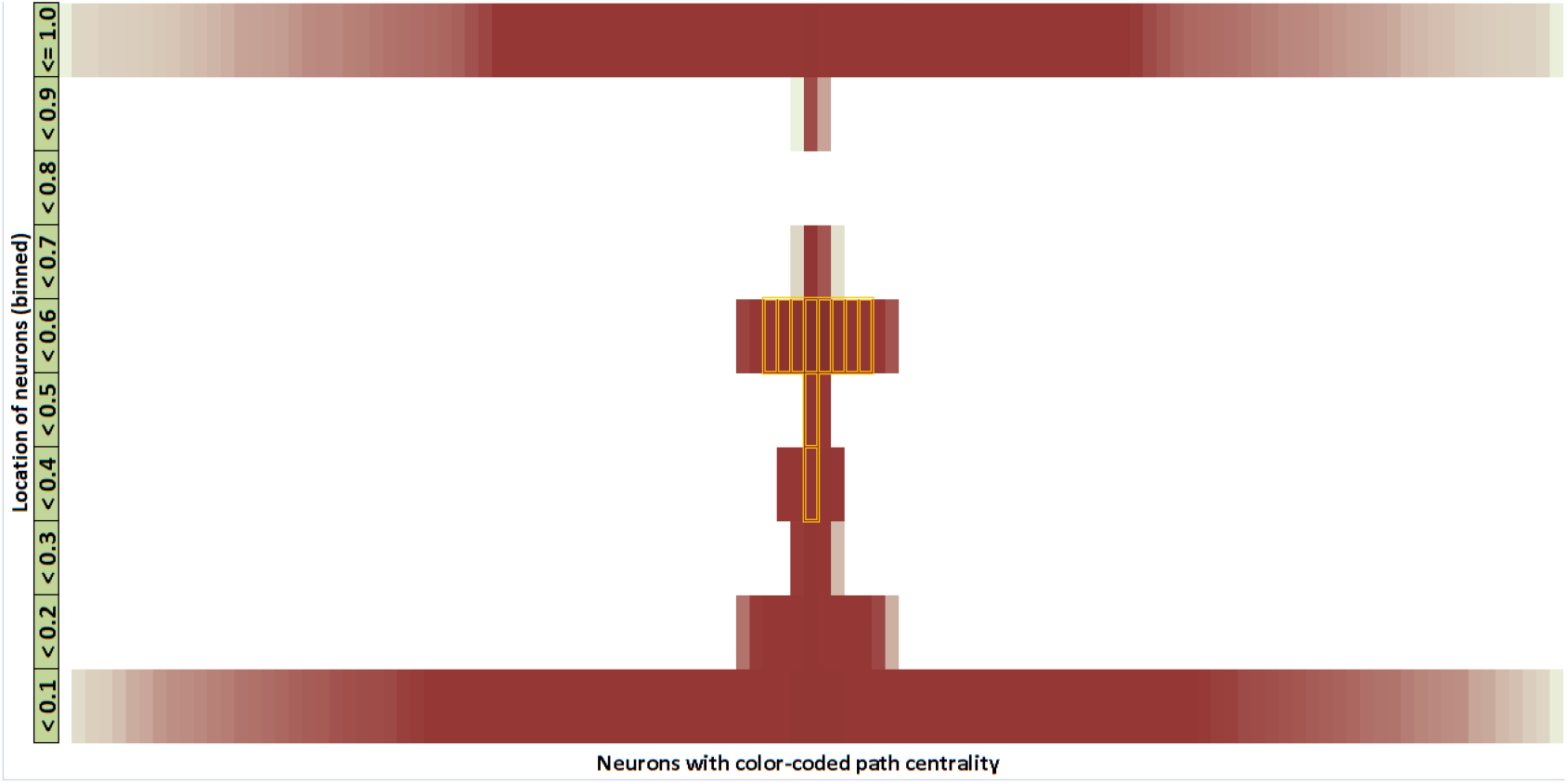
Visualization of the *C. elegans* connectome based on the location metric. We discretize the location metric in 10 bins (each bin accounting for 1/10 of the 0 − 1 range). The path centrality of each node is represented by its color intensity (darker for higher path centrality). Nodes with higher centrality are placed closer to the vertical mid-line. The core nodes are marked with an orange outline. The location of all core neurons falls between 0.3-0.6, close to the middle of the hourglass.

### *C. elegans* as an Encoder-Decoder Architecture

We can think of *C. elegans* as an information processing system that transforms input information, collected and encoded by sensory neurons, to output information that is represented by the activity of motor neurons. The analysis of the previous sections has identified a number of *core neurons* that most of the sensory-to-motor neural pathways go through. The exact number of core neurons depends on the fraction *τ* of all sensory-to-motor paths covered by the core.

Suppose that a given set of core neurons forms the intermediate set *Z*, defined in Section “Encoder-Decoder Architecture”. We can then compute the number of paths *P*_*S*_ (*Z*) from the set *S* of all sensory neurons to the neurons in *Z* as a proxy for the information processing cost of an encoding operation that transforms *S* to *Z*. Similarly, the number of paths *P*_*S*_ (*Z*) from the neurons in *Z* to the set *T* of all motor neurons can be thought of as a proxy for the information processing cost of a decoding operation that transforms *Z* to *T*. We also need to consider any sensory-to-motor paths *P_S,b_*(*T*) that bypass the core neurons in *Z* — this is a proxy for the cost of any additional information processing that is specific to each motor neuron and that is not provided by the encoding-decoding function of *Z*.

These three cost terms are shown in Figure 12 as we increase the number of core neurons included in *Z* (i.e., as we increase the threshold *τ*). The bypass-*Z* cost is the dominant cost term until we include about 15 neurons in *Z*. This suggests that the information provided by sensory neurons cannot be captured well with fewer neurons. On the other hand, the costs of the encoding and decoding operations (*P*_*S*_ (*Z*) and *P*_*S*_ (*Z*), respectively) increase with the number of neurons in *Z*, as expected.

**Figure 12:**
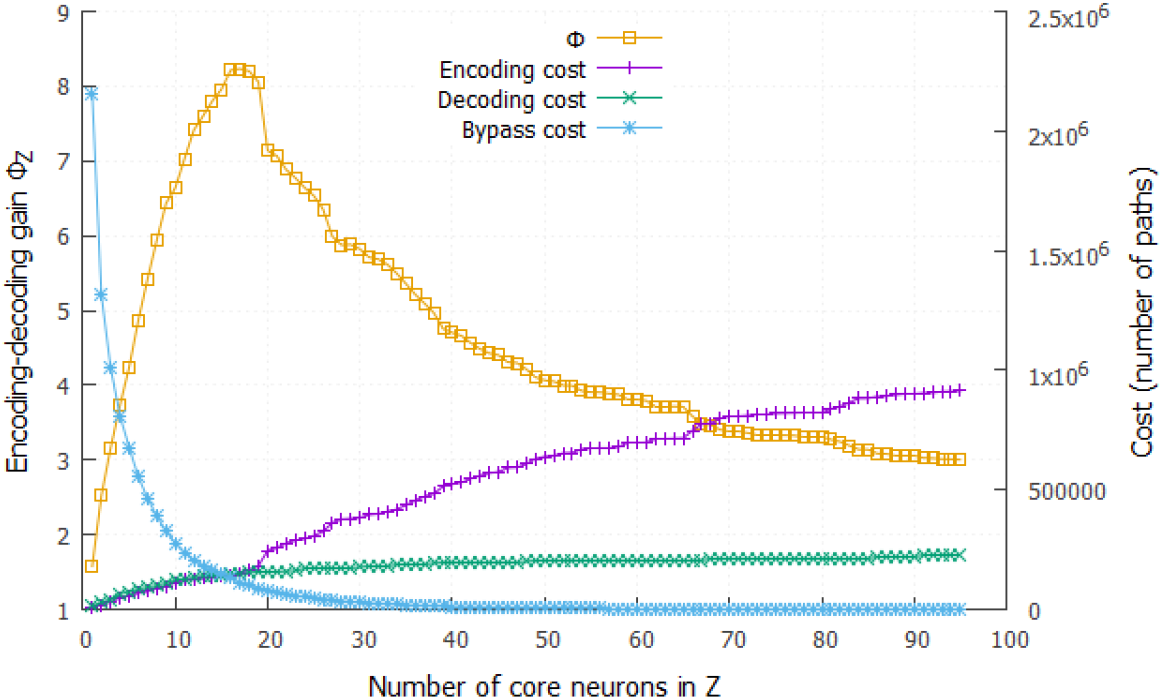
The encoder-decoder gain ratio Φ_*Z*_ as the number of core neurons in the encoding set *Z* increases (yellow curve). The maximum value of Φ_*Z*_ is about 8.2 when *Z* includes the first 16 core neurons. Based on the cost framework of Section “Encoder-Decoder Architecture”, this means that the hourglass organization of the *C. elegans* connectome reduces the sensory-to-motor information processing cost eight-fold. The figure also shows the three relevant cost terms: cost of encoding the information provided by sensory neuron using neurons in *Z* (magenta), cost of decoding that information to drive all motor neurons (green), and cost of processing pathways that bypass the core (blue).

The encoder-decoder gain ratio Φ_*Z*_ (see Equation 6) shows that the maximum cost reduction takes place when we consider the first 16 core neurons (corresponds to *τ* =95% for the *SP* ^+2^ set of paths). In that case, the encoder-decoder architecture achieves an eight-fold decrease (Φ_*Z*_ = 8.2) in terms of information processing cost relative to a hypothetical architecture in which the information processing cost of each motor neuron is computed separately, based on the number of paths from sensory neurons to that motor neuron.

An important question is whether the hourglass architecture achieves this cost reduction by increasing the path length between sensory and motor neurons (in terms of the number of neurons in each path). This trade-off between *network efficiency* (associated with the distribution of path lengths in a network) and *network cost* has received significant attention in network neuroscience [48, 57, 49]. Networks that minimize the length of every processing path connect every source to every target with a direct link – a costly design approach. On the other hand, networks that attempt to reduce the number of intermediate links typically need longer source-to-target paths (for the same reason that flying between two cities often requires one or more intermediate stops).

Here, we examine whether the hourglass architecture introduces a significant increase in the average path length from sensory to motor neurons relative to the ensemble of randomized networks. Recall that those networks do not follow the hourglass architecture (see Figure 9) but they maintain the in-degree of each neuron and the hierarchical ordering between neurons. Given that each neuron selects randomly its inputs from any neuron that is “lower” in the hierarchy (closer to the sensory neurons), we expect that such randomized non-hourglass networks will be more efficient (i.e., they will have shorter paths). In the extreme case that every motor neuron receives connections only from sensory neurons, the average path length will be minimized.

Figure 13 shows that the randomized networks are not as cost-efficient as the original *C. elegans* connectome, (their encoder-decoder gain ratio is around 2 even though the original network’s is 8.2). However, the randomized networks provide shorter path lengths. Their average sensory-to-motor path length is 3.9, according to the *SP* ^+2^ set of paths, while the same metric for the original connectome is 5.4 hops. In other words, the hourglass organization of the *C. elegans* connectome trades off the reuse of intermediate-level neurons and connections with a modest increase in the length of sensory-to-motor path lengths. We should mention that this reduction in path-length efficiency is smaller for other path sets; for instance, with the set of shortest paths (*SP*) the average path length of the original connectome is 3.4 hops while the randomized connectomes have a mean of 3.0.

**Figure 13:**
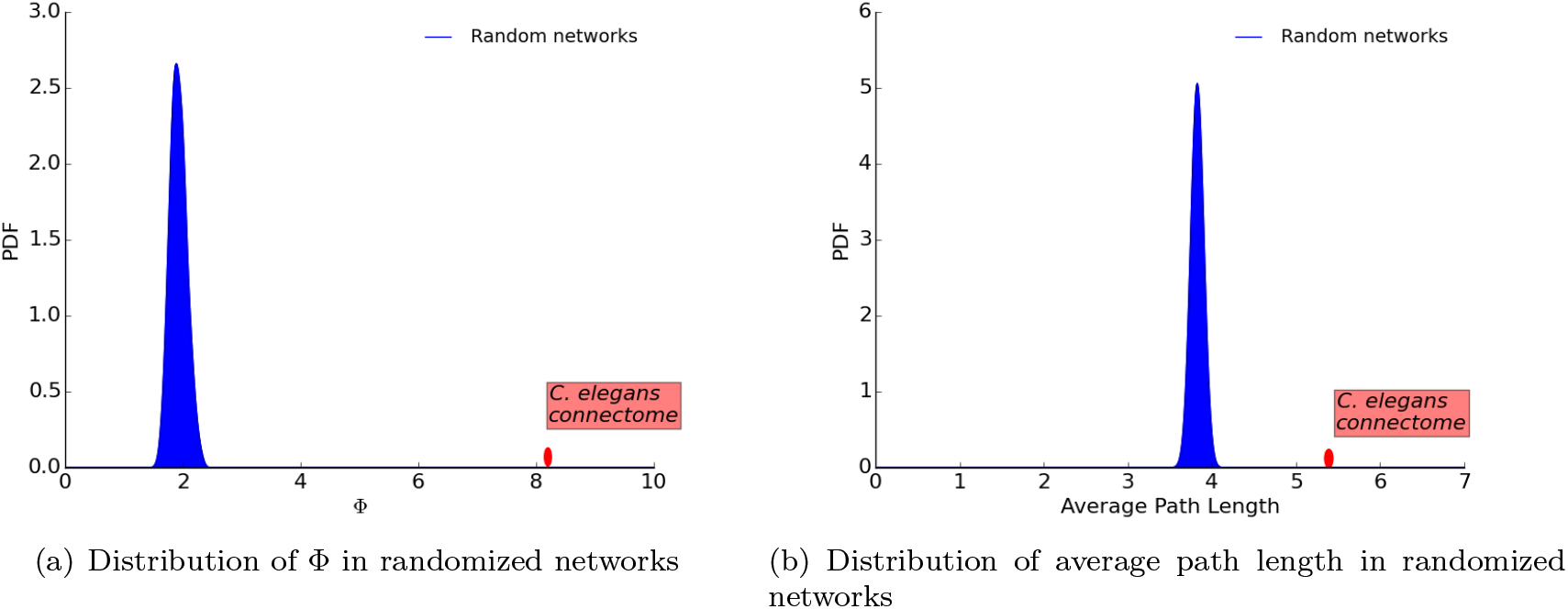
The ensemble of randomized networks have much lower encoding-decoding gain Φ than the original *C. elegans* connectome (a) – but not significantly lower average path length (b).

### Including Gap Junctions: the Complete Network

Gap junctions provide a different type of connectivity between neurons than chemical synapses. Chemical synapses use neurotransmitters to transfer information from the presynaptic to the post-synaptic neuron, while gap junctions work by creating undirected electrical channels and they provide faster (but typically weaker) neuronal coupling. The directionality of the current flow in gap junctions cannot be detected from electron micrographs. Hence they are treated as bidirectional in the *C. elegans* connectome.

In this section, we consider both gap junctions and chemical synapses, forming what we refer to as the *complete network*. Gap junctions connect 253 (out of 279) neurons through 514 undirected connections (that we treat as 1028 directional connections). Recall that the number of (directed) chemical connections is 2194. 64% of the gap junction connections do not co-occur with chemical connections between the same pair of neurons, while 83% of the synaptic connections do not co-occur with gap junction connections. In other words, the inclusion of gap junctions changes significantly the connectivity between neurons.

The complete network has 1180 FF, 1468 LT and 574 FB connections. If we remove feedback connections, as we did for the synaptic network, we end up with a total of 2648 directed connections. The inclusion of gap junctions also increases significantly the number of paths between sensory and motor neurons, independent of the routing method. If we focus on *SP* ^+2^, the number of paths increases by a factor of 2.3 (about 7.7 millions).

Figure 14(a) shows the cumulative path coverage as a function of the number of nodes in the core. Figure 14(b) examines the effect of the path coverage threshold *τ* on the resulting H-score. Both curves are quite similar to the corresponding results for the synaptic network.

**Figure 14:**
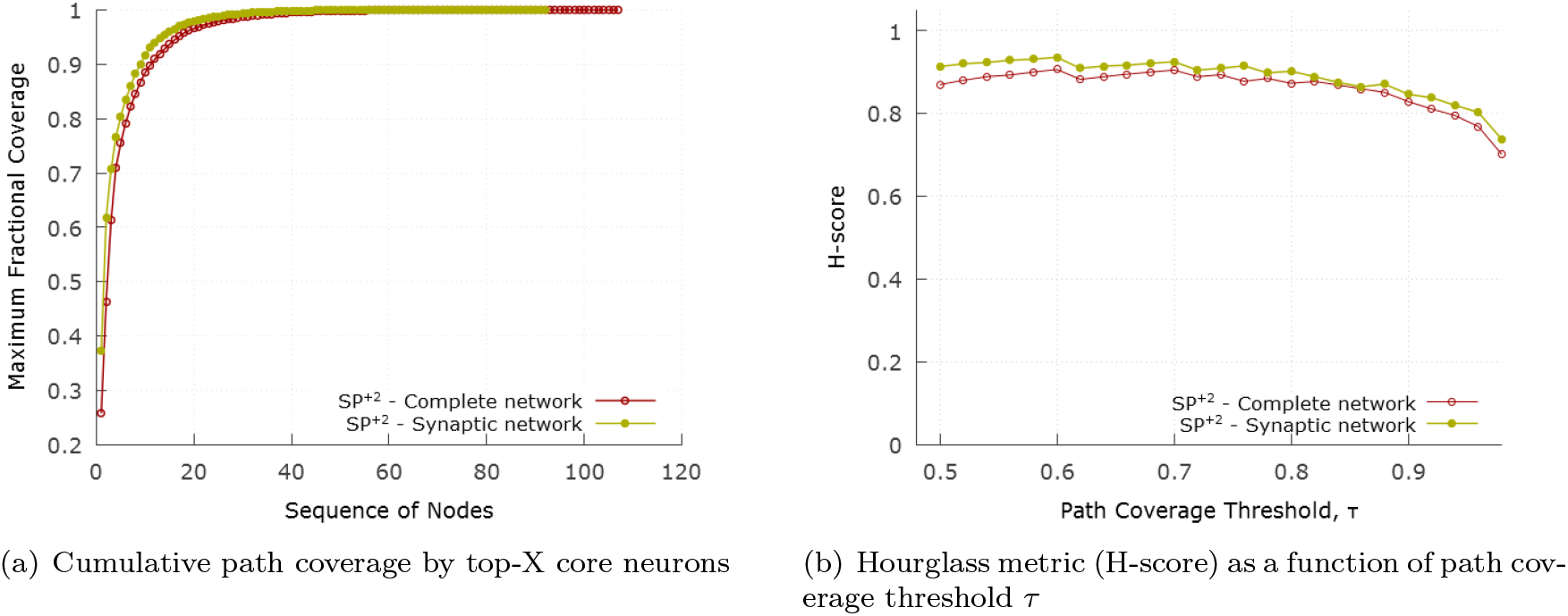
Hourglass analysis for complete network with the *SP* ^+2^ set of paths. (a) compares the cumulative path coverage of top core neurons between synaptic and complete network. (b) compares the progression of H-score between synaptic and complete network as *τ* varies.

With *τ* = 90%, the resulting core nodes are shown in Table 3. The H-score for the complete network is 0.83 (compared to 0.87 for the synaptic network).

**Table 3:**
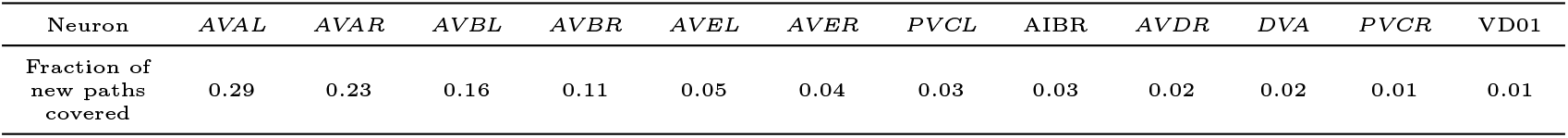
The identified 12 core neurons in the complete network. The 10 neurons shown in italic were also the core of the synaptic network.

| Neuron | *AVAL* | *AVAR* | *AVBL* | *AVBR* | *AVEL* | *AVER* | *PVCL* | *AIBR* | *AVDR* | *DVA* | *PVCR* | *VD01* |
| --- | --- | --- | --- | --- | --- | --- | --- | --- | --- | --- | --- | --- |
| Fraction of new paths covered | 0.29 | 0.23 | 0.16 | 0.11 | 0.05 | 0.04 | 0.03 | 0.03 | 0.02 | 0.02 | 0.01 | 0.01 |

The two additional core neurons that appear in the hourglass waist of the complete network but not in the synaptic network are:

- AIBR: related to locomotion, food and odor evoked behaviors, local search, lifespan and starvation response.
- VD01: related to motor-Sinusoidal body movement-locomotion.

The encoder-decoder gain analysis for the complete network appears in Figure S4. Qualitatively the encoder-decoder gain ratio follows the same trend with the network of only chemical synapses (see Figure 12) but the maximum value of Φ_*Z*_ is slightly less (7.4 instead of 8.2).

## Discussion

In this Section, we discuss in more detail prior studies that relate to the hourglass effect in *C. elegans* or more broadly in neuroscience. Varshney et al. [43] analyzed the structural properties of the *C. elegans* connectome and found that several central neurons (based on closeness centrality) play a key role in information processing. Among them are command inter-neurons such as *AVA, AVB, AVE* that are responsible for locomotion control. On the other hand, neurons such as *DVA* or *ADE* have high out-closeness centrality and a good position to propagate a signal to the rest of the network. Most of the “central” neurons in that study are also included in the hourglass core.

The modular organization of the *C. elegans* connectome has been discovered by Sohn et al. [58] through cluster analysis. Their analysis showed that communities correspond well to known functional circuits and it helped uncover the role of a few previously unknown neurons. They also identified a hierarchical organization among five key clusters that form a backbone for higher-order complex behaviors.

The fact that the rich-club interneurons are almost identical with the hourglass core, even though these two network properties are qualitatively different, suggests that these 10-15 neurons form a information processing bottleneck that does not simply compress but also *integrates* the information from different sensory modalities, before driving any higher-level neural circuits.

This hypothesis is also supported by the analysis of functional modules in the *C. elegans* connectome, by Pan et al. [59], which showed that neurons in the same module are located close and contribute in the same task. That study identified 23 *connector hub* neurons, i.e., high-connectivity neurons that connect to most or all functional modules. The eleven core neurons that we identified with the *SP* ^+2^ paths also belong in that set of connector hubs. The fact that all hourglass core neurons are also connector hubs between functional modules supports the idea that these neurons integrate multimodal information, rather than simply compress the sensory information in a segregated manner. Note that the distinction between connector hubs, non-hub connectors, etc, depends on certain thresholds and so it is not surprising that some connector hubs such as AVKL or SMBVL do not appear in the hourglass core.

The posterior nervous system of the male *C. elegans* connectome was analyzed by Jarrell et al. [60] (recall that we analyze the hermaphrodite *C. elegans* connectome). One of their conclusions was that the nervous system has a mostly feedforward architecture that runs from sensory to motor neurons via interneurons. There is also some feedback circuitry in the nervous system and the actual physical output of the worm (i.e. motion etc.) feeds back to sensory neurons to allow closed-loop control. There are however many more feedforward loops (termed lateral connections in our analysis) that provide localized coordination most notably visible within interneurons. More recently, the same research group has mapped the complete connectome of the male nematode, focusing on its differences with the hermaphrodite [61].

Yan et al. have applied a controllability framework to analyze the *C. elegans* connectome, aiming to identify essential neurons for locomotion [62]. Some of those neurons also appear in the hourglass core (AVAL/R, AVBL/R, AVDL/R, PVCL/R) – but there are also several neurons (such as the six neurons of the DD class) that do not stand out in the hourglass analysis. This is not surprising given that the two studies ask very different questions: Yan et al. ask which neurons are essential to control every motor neuron or muscle, while we ask which neurons form a bottleneck in the feedforward flow of information from sensory to motor neurons.

The physical placement of neurons in *C. elegans* has been thought to be not exclusively optimized for global minimum wiring but rather for a variety of other factors of which the minimization of pair-wise processing steps is important. For example, Kaiser and Hilgetag [49] showed that the total wiring length can be reduced by 48% by optimally placing the neurons. However that would significantly increase the number of processing nodes along shortest paths between components as well. Similar findings were also revealed by Chen et al. [57], concluding that the placement of neurons does not globally minimize wiring length. These studies emphasize the notion of choosing shorter communication paths between neuron pairs and supports our approach of choosing paths that are shortest, or close to shortest, in terms of processing steps.

Analysis by Csoma et al. [63] challenged the well rooted notion of shortest path based communication routing in the human brain network. They collected empirical data through diffusion MRI and concluded that although a large number of paths conform to the shortest path assumption, a significant fraction (20-40%) are inflated up to 4-5 hops.

Research by Avena-Koenigsberger et al. [47, 64] analyzed in depth the communication strategies in the human brain and also challenged the shortest path assumption. They discussed how the computation of shortest path routing is not feasible in the brain circuitry, and the shortest path routes would leave out around 80% of neural connections. They examined the spectrum of routing strategies hinging upon the amount of global information and communication required. At one end of the spectrum, there are random-walk routing mechanisms that are wasteful and often fail to achieve efficient routes but require no knowledge. On the other end there is shortest-path routing requiring global wiring knowledge at each neuron. As a more realistic choice, they studied the *k*-shortest path based approach (with *k* being 100). Their findings show that this strategy increases the utilization of connections. We have used a more relaxed constraint to choose paths between any two nodes by allowing all possible paths that are up to 2 hops longer than the shortest path between the corresponding pair.

Markov et al. have shown that the macaque cortical network includes a highly interconnected “bow-tie core” [42]. At first, this may seem relevant to the hourglass effect. We should note however that the network of Markov et al. considers 29 cortical regions and 17 of them are in the bow-tie core. On the contrary, a defining characteristic of the hourglass effect is that the number of core nodes at the waist is a small fraction of the total network size.

In some earlier studies, the hourglass effect is defined for layered networks, based on on the number of nodes at each layer. A network is referred to as an hourglass if the width of the intermediate layers is much smaller relative to the width of the input and output layers [24, 35, 65]. In this work, we generalize the definition of the hourglass effect to include networks that do not have clearly defined layers and that include feedback or lateral connections.

What is the biological significance of the hourglass architecture in the *C. elegans* connectome? Is it just an interesting graph-theoretic property or does this architecture provide an adaptive advantage that could be selected by evolution?

First, it is important to set appropriate expectations for any study that analyzes the connectome attempting to learn something valuable about the underlying biology. It has been argued by several authors, including C. Bargmann and E. Marder [66], that mechanisms such as neuromodulators, parallel and antagonistic pathways and circuits, and complex neuronal dynamics can completely change the function of a given neural circuit. We believe that a connectome should be viewed as an *architectural constraint that limits the scope of possible functions that a neural circuit can perform* – rather than as the unique determinant of those functions.

The earlier *C. elegans* literature has attributed specific functions to the “command interneurons” or it has associated those interneurons with one or more functional circuits (for instance, see [67, 68]). The main contribution of our study is to propose a different way to think about the role of those interneurons in the *C. elegans* connectome: the interneurons between sensory and motor neurons can be thought of as forming an *encoder-decoder network*. This network reduces the intrinsic dimensionality of the low-level sensory information, and then integrates the compressed information from different sensory modalities to compute few intermediate-level sub-functions. The latter are then combined and re-used in higher-level behavioral circuits and tasks. Those few sub-functions are encoded in the activity of 10-15 core interneurons in the hourglass waist.

So, instead of trying to identify the function of each neuron in the connectome, or instead of focusing on individual functional circuits ignoring all others behaviors and circuits, we can focus on that smaller set of 10-15 core interneurons and attempt, through a combination of experiments and modeling, to reverse engineer the sub-functions they “compute.” These sub-functions will probably be much simpler than the observable behaviors of the organism (e.g., escape response or social feeding) – they can be viewed as *re-usable functional modules.* Then, for each of the observable behaviors of the organism, we can try to find out how that task is accomplished by combining in different ways those functional modules. We firmly believe that such a research agenda will be more tractable because it depends on a smaller number of components (10-15) that need to be “reverse engineered”, compared to the number of all neurons in *C. elegans*.

The core neurons at the hourglass waist create a “bottleneck” in the flow of information from sensory to motor neurons. Such bottleneck effects have been studied in the literature under different names. The most relevant such framework is the *information bottleneck* method developed based on information theory results: given a joint probability distribution between an input vector *X* and an output vector *Y*, the goal of that method is to compute an optimal intermediate-level representation *T* that is both compact (i.e., a highly compressed version of *X*) and able to predict *Y* accurately [69, 70]. It appears that the *C. elegans* connectome has evolved to “compute” such a compact and integrated intermediate-level representation of its sensory inputs, represented by the 10-15 core interneurons at the hourglass waist.

## Supplementary Information

**Figure S1:**
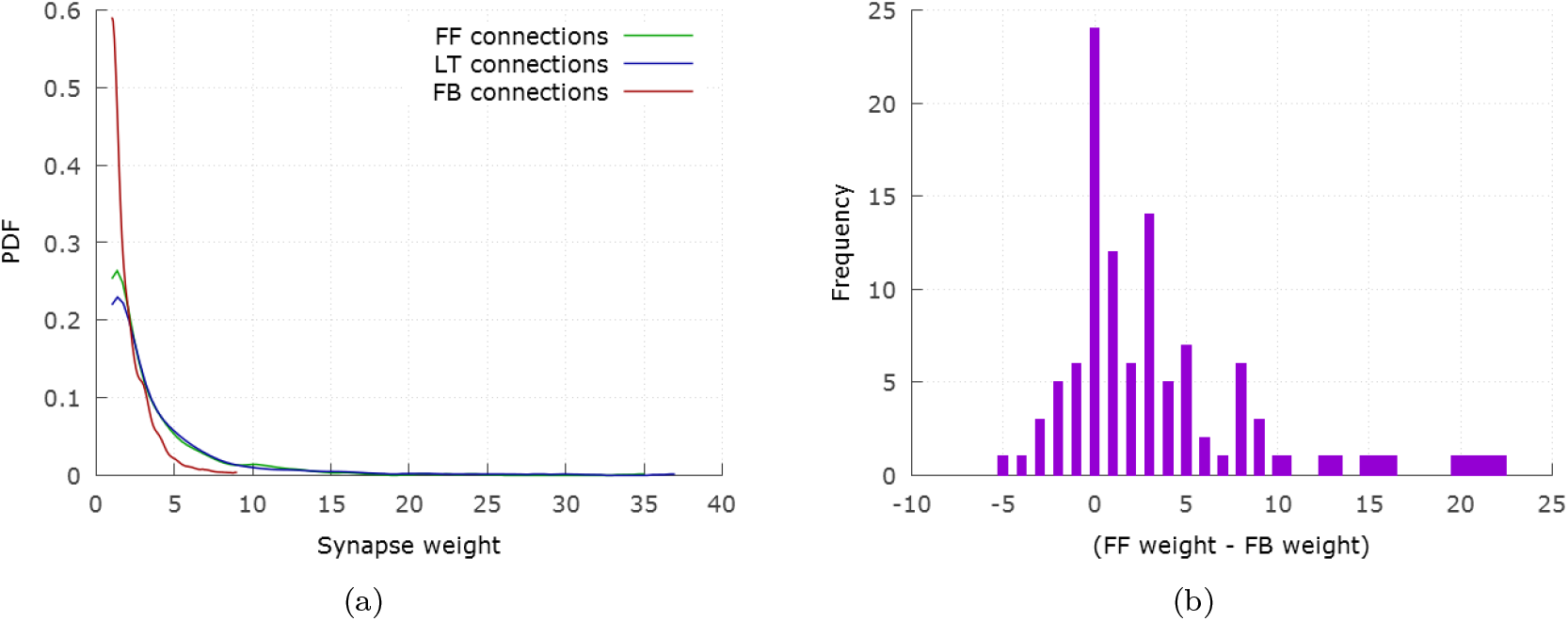
(a) Weight distribution of FF, LT and FB connections. (b) Considering only pairs of neurons with reciprocal FF and FB connections, this histogram shows the difference of the FF weight minus the FB weight.

**Figure S2:**
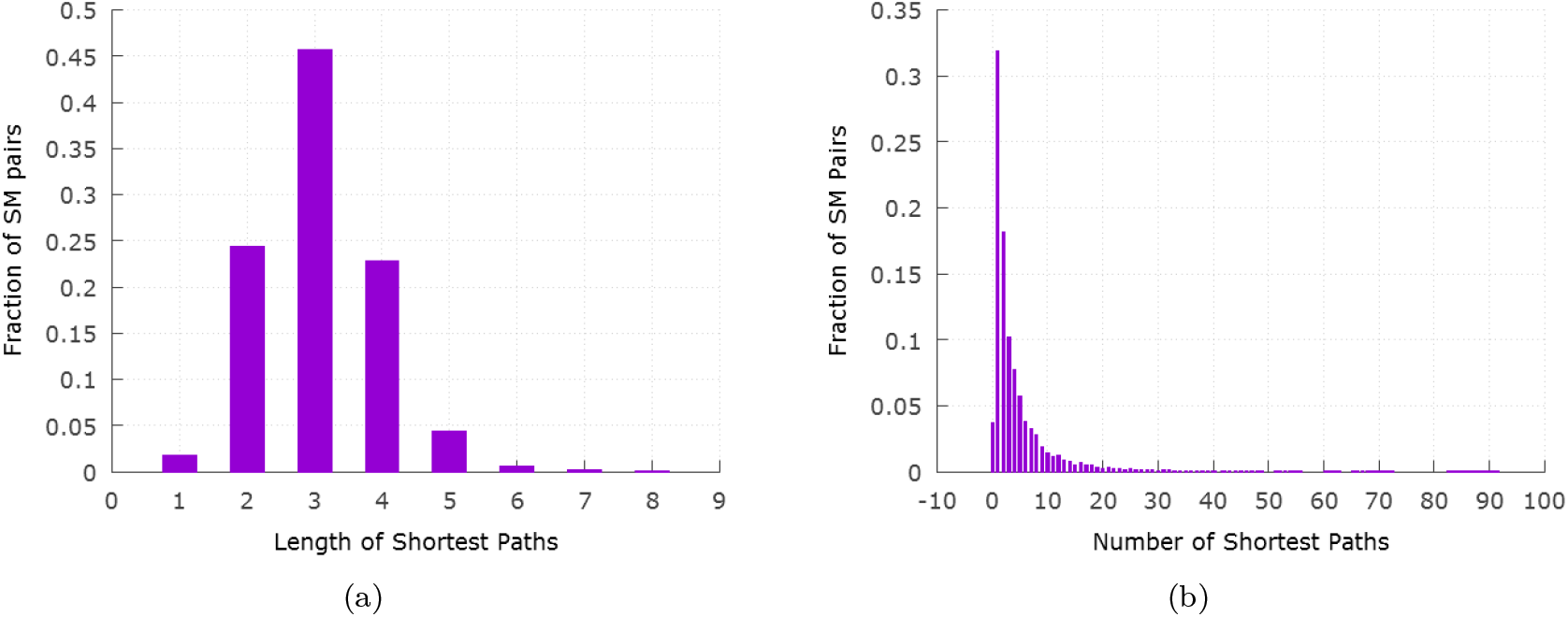
(a) The length distribution for all shortest paths from sensory to motor neurons. Almost all shortest paths are shorter than 6 hops. (b) Distribution of the number of distinct shortest paths from a sensory neuron to a motor neuron. For about 50% of S-M pairs, there are more than two shortest paths.

**Figure S3:**
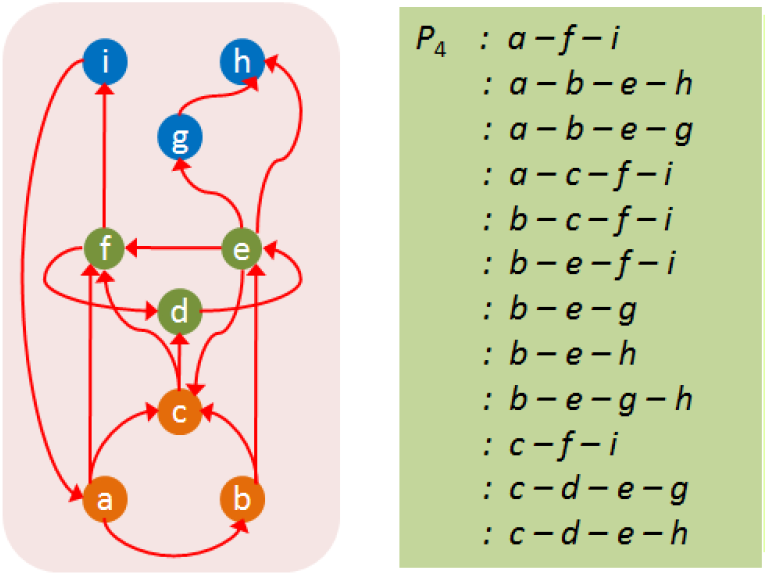
All paths for the routing scheme *P*_4_. The model network is the same one depicted in Figure 3.

**Table S1:** In the synaptic network, the top-5% strongest connections are dominated by FF connections from S neurons to I or M neurons, and by LT connections between I neurons and M neurons. On the other hand, none of the FB connections appear in this set.

| Connection type | Connection sub-type | Fraction in: |  |
| --- | --- | --- | --- |
|  |  | entire connectome | top-5% weighted connections |
| FF | S-I | 0.18 | 0.35 |
|  | S-M | 0.08 | 0.02 |
|  | I-M | 0.16 | 0.15 |
| LT | S-S | 0.09 | 0.00 |
|  | I-I | 0.22 | 0.15 |
|  | M-M | 0.14 | 0.33 |
| FB | I-S | 0.06 | 0.00 |
|  | M-I | 0.06 | 0.00 |
|  | M-S | 0.01 | 0.00 |

**Table S2:**
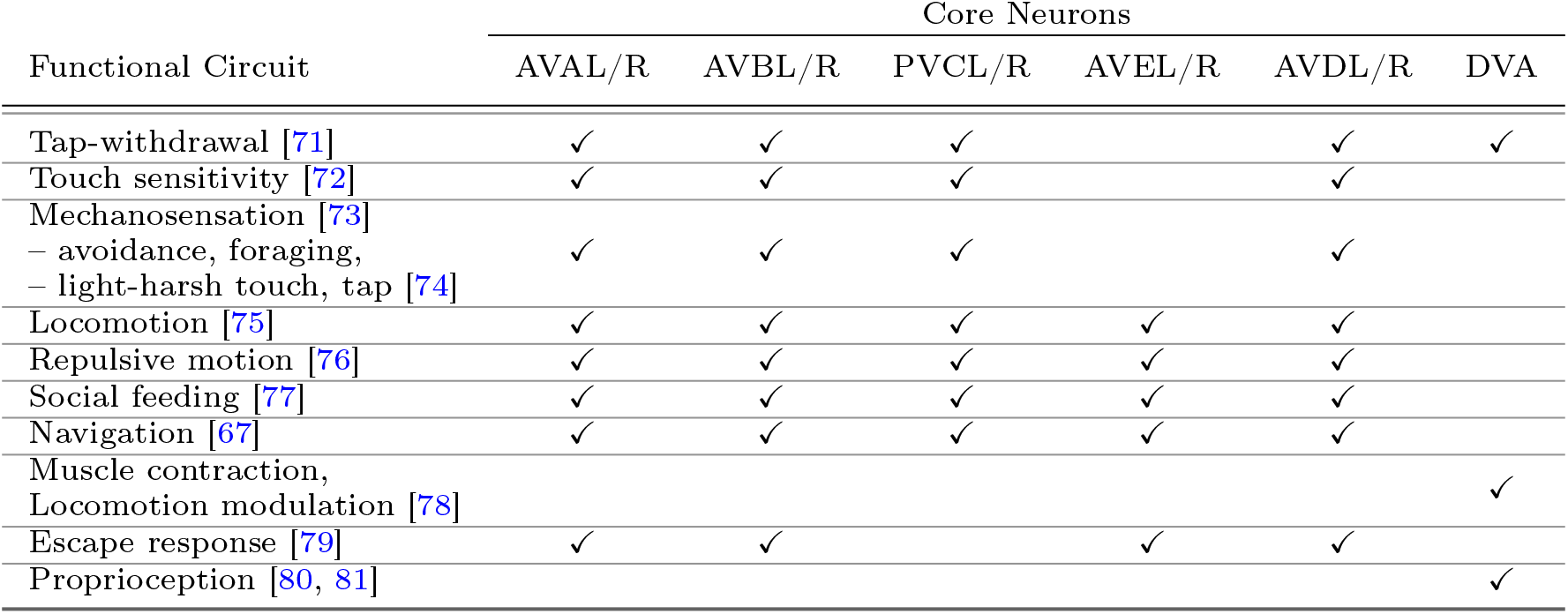
Functional circuits associated with core neurons based on the *C. elegans* literature. The core neurons appear in several circuits, mostly related to spontaneous or planned movement. Many of the adaptive behaviors of the organism such as feeding, egg-laying, escape and navigation require a common set of underlying simpler tasks. Some of the circuits shown (e.g. thermotaxis, chemosensation, olfactory behavior) perform tasks that start with activity in some sensory neurons, followed by a locomotory response that is modulated by certain core interneurons.

**Figure S4:**
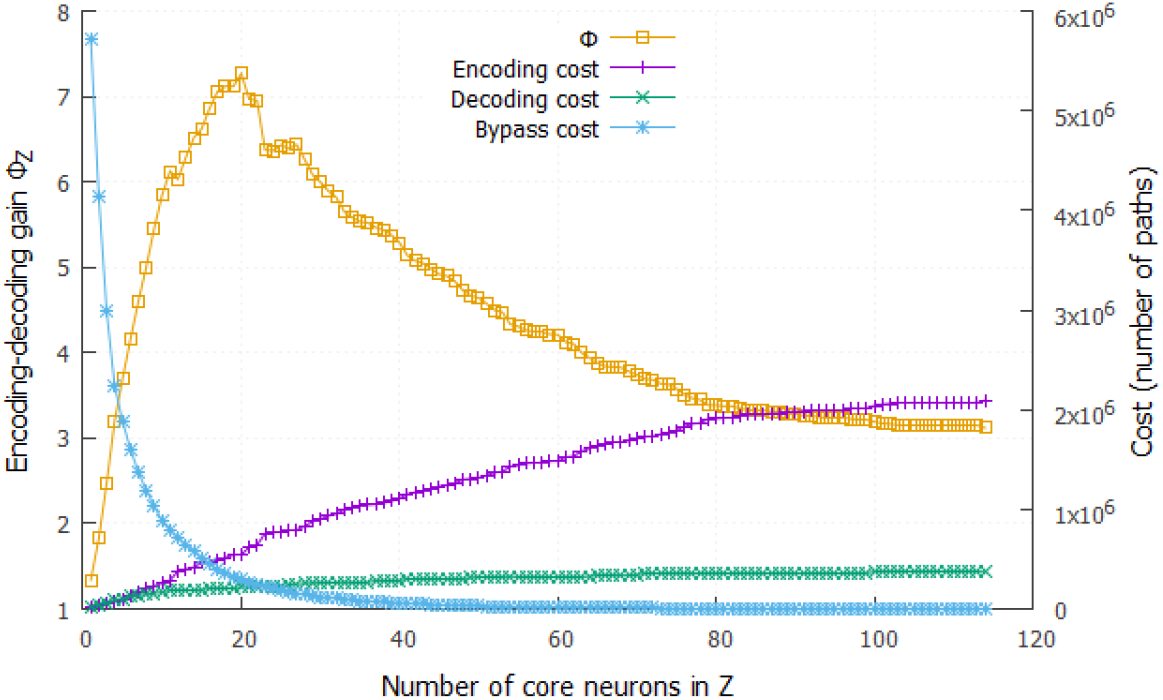
The encoder-decoder gain ratio Φ_*Z*_ for the combined network containing both chemical synapses and gap junctions (contrast with Figure 12). The maximum value of Φ_*Z*_ is 7.4 when *Z* includes the first 20 core neurons. Recall that the maximum value of Φ_*Z*_ in the network of chemical synapses is 8.2.

## References

1. Bassett DS, Gazzaniga MS. Understanding complexity in the human brain. Trends in cognitive sciences. 2011;15(5):200–209.

2. Meunier D, Lambiotte R, Bullmore ET. Modular and hierarchically modular organization of brain networks. Frontiers in Neuroscience. 2010;4:200.

3. Parnas DL, Clements PC, Weiss DM. The modular structure of complex systems. In: Proceedings of the 7th International Conference on Software Engineering. IEEE Press; 1984. p. 408–417.

4. Schilling MA. Toward a general modular systems theory and its application to interfirm product modularity. Academy of Management Review. 2000;25(2):312–334.

5. Baldwin CY, Clark KB. Design Rules: The Power of Modularity. vol. 1. MIT press, Cambridge; 2000.

6. Callebaut W, Rasskin-Gutman D. Modularity: Understanding the Development and Evolution of Natural Complex Systems. MIT press, Cambridge; 2005.

7. Wagner GP, Pavlicev M, Cheverud JM. The road to modularity. Nature Reviews Genetics. 2007;8(12):921–931.

8. Ravasz E, Barabási AL. Hierarchical organization in complex networks. Physical Review E. 2003;67(2):026112.

9. Sales-Pardo M, Guimera R, Moreira AA, Amaral LAN. Extracting the hierarchical organization of complex systems. Proceedings of the National Academy of Sciences. 2007;104(39):15224–15229.

10. Simon HA. The architecture of complexity. Springer; 1991.

11. Yu H, Gerstein M. Genomic analysis of the hierarchical structure of regulatory networks. Proceedings of the National Academy of Sciences. 2006;103(40):14724–14731.

12. Clune J, Mouret JB, Lipson H. The evolutionary origins of modularity. Proceedings of the Royal Society of London B: Biological Sciences. 2013;280(1755):20122863.

13. Mengistu H, Huizinga J, Mouret JB, Clune J. The evolutionary origins of hierarchy. PLoS computational biology. 2016;12(6):e1004829.

14. Fortuna MA, Bonachela JA, Levin SA. Evolution of a modular software network. Proceedings of the National Academy of Sciences. 2011;108(50):19985–19989.

15. Huang CC, Kusiak A. Modularity in design of products and systems. Systems, Man and Cybernetics, Part A: Systems and Humans, IEEE Transactions on. 1998;28(1):66–77.

16. Myers CR. Software systems as complex networks: Structure, function, and evolvability of software collaboration graphs. Physical Review E. 2003;68(4):046116.

17. Kashtan N, Alon U. Spontaneous evolution of modularity and network motifs. Proceedings of the National Academy of Sciences of the United States of America. 2005;102(39):13773–13778.

18. Kashtan N, Noor E, Alon U. Varying environments can speed up evolution. Proceedings of the National Academy of Sciences. 2007;104(34):13711–13716.

19. Lorenz DM, Jeng A, Deem MW. The emergence of modularity in biological systems. Physics of Life Reviews. 2011;8(2):129–160.

20. Kirsten H, Hogeweg P. Evolution of networks for body plan patterning; interplay of modularity, robustness and evolvability. PLoS Comput Biol. 2011;7(10):e1002208.

21. Kitano H. Biological robustness. Nature Reviews Genetics. 2004;5(11):826–837.

22. Stelling J, Sauer U, Szallasi Z, Doyle FJ, Doyle J. Robustness of cellular functions. Cell. 2004;118(6):675–685.

23. Sabrin KM, Dovrolis C. The hourglass effect in hierarchical dependency networks. Network Science. 2017;5(4):490–528.

24. Friedlander T, Mayo AE, Tlusty T, Alon U. Evolution of bow-tie architectures in biology. PLoS computational biology. 2015;11(3):e1004055.

25. Casci T. Development: Hourglass theory gets molecular approval. Nature Reviews Genetics. 2011;12(2):76–76.

26. Quint M, Drost HG, Gabel A, Ullrich KK, Bönn M, Grosse I. A transcriptomic hourglass in plant embryogenesis. Nature. 2012;490(7418):98–101.

27. Tanaka R, Csete M, Doyle J. Highly optimised global organisation of metabolic networks. IEE Proceedings-Systems Biology. 2005;152(4):179–184.

28. Zhao J, Yu H, Luo JH, Cao ZW, Li YX. Hierarchical modularity of nested bow-ties in metabolic networks. BMC Bioinformatics. 2006;7(1):386.

29. Beutler B. Inferences, questions and possibilities in Toll-like receptor signalling. Nature. 2004;430(6996):257–263.

30. Oda K, Kitano H. A comprehensive map of the toll-like receptor signaling network. Molecular Systems Biology. 2006;2(1).

31. Supper J, Spangenberg L, Planatscher H, Dräger A, Schröder A, Zell A. BowTieBuilder: modeling signal transduction pathways. BMC Systems Biology. 2009;3(1):1.

32. Quiroga RQ, Reddy L, Kreiman G, Koch C, Fried I. Invariant visual representation by single neurons in the human brain. Nature. 2005;435(7045):1102–1107.

33. Riesenhuber M, Poggio T. Hierarchical models of object recognition in cortex. Nature Neuroscience. 1999;2(11):1019–1025.

34. Hinton GE, Salakhutdinov RR. Reducing the dimensionality of data with neural networks. Science. 2006;313(5786):504–507.

35. Akhshabi S, Dovrolis C. The evolution of layered protocol stacks leads to an hourglass-shaped architecture. ACM SIGCOMM Computer Communication Review. 2011;41(4):206–217.

36. Swaminathan JM, Smith SF, Sadeh NM. Modeling supply chain dynamics: A multiagent approach*. Decision Sciences. 1998;29(3):607–632.

37. Csermely P, London A, Wu LY, Uzzi B. Structure and dynamics of core/periphery networks. Journal of Complex Networks. 2013;1(2):93–123.

38. Holme P. Core-periphery organization of complex networks. Physical Review E. 2005;72(4):046111.

39. Csete M, Doyle J. Bow ties, metabolism and disease. TRENDS in Biotechnology. 2004;22(9):446–450.

40. Domazet-Lošo T, Tautz D. A phylogenetically based transcriptome age index mirrors ontogenetic divergence patterns. Nature. 2010;468(7325):815–818.

41. Ma HW, Zeng AP. The connectivity structure, giant strong component and centrality of metabolic networks. Bioinformatics. 2003;19(11):1423–1430.

42. Markov NT, Ercsey-Ravasz M, Van Essen DC, Knoblauch K, Toroczkai Z, Kennedy H. Cortical high-density counterstream architectures. Science. 2013;342(6158):1238406.

43. Varshney LR, Chen BL, Paniagua E, Hall DH, Chklovskii DB. Structural properties of the *Caenorhabditis elegans* neuronal network. PLoS computational biology. 2011;7(2):e1001066.

44. Towlson EK, Vértes PE, Ahnert SE, Schafer WR, Bullmore ET. The rich club of the *C. elegans* neuronal connectome. Journal of Neuroscience. 2013;33(15):6380–6387.

45. Goodfellow I, Bengio Y, Courville A. Deep learning. MIT press; 2016.

46. Reece JB, Urry LA, Cain ML, Wasserman SA, Minorsky PV, Jackson RB, et al. Campbell biology. Pearson Boston; 2014.

47. Avena-Koenigsberger A, Misic B, Sporns O. Communication dynamics in complex brain networks. Nature Reviews Neuroscience. 2018;19(1):17.

48. Bullmore E, Sporns O. The economy of brain network organization. Nature Reviews Neuroscience. 2012;13(5):336.

49. Kaiser M, Hilgetag CC. Nonoptimal component placement, but short processing paths, due to long-distance projections in neural systems. PLoS computational biology. 2006;2(7):e95.

50. Raj A, Chen Yh. The wiring economy principle: connectivity determines anatomy in the human brain. PloS one. 2011;6(9):e14832.

51. Abdelnour F, Voss HU, Raj A. Network diffusion accurately models the relationship between structural and functional brain connectivity networks. Neuroimage. 2014;90:335–347.

52. Mišić B, Betzel RF, Nematzadeh A, Goni J, Griffa A, Hagmann P, et al. Cooperative and competitive spreading dynamics on the human connectome. Neuron. 2015;86(6):1518–1529.

53. Ishakian V, Erdös D, Terzi E, Bestavros A. A Framework for the Evaluation and Management of Network Centrality. In: SDM. SIAM; 2012. p. 427–438.

54. Karrer B, Newman ME. Random graph models for directed acyclic networks. Physical Review E. 2009;80(4):046110.

55. Colizza V, Flammini A, Serrano MA, Vespignani A. Detecting rich-club ordering in complex networks. Nature physics. 2006;2(2):110.

56. Hagberg A, Swart P, S Chult D. Exploring network structure, dynamics, and function using NetworkX. Los Alamos National Lab.(LANL), Los Alamos, NM (United States); 2008.

57. Chen BL, Hall DH, Chklovskii DB. Wiring optimization can relate neuronal structure and function. Proceedings of the National Academy of Sciences. 2006;103(12):4723–4728.

58. Sohn Y, Choi MK, Ahn YY, Lee J, Jeong J. Topological cluster analysis reveals the systemic organization of the *Caenorhabditis elegans* connectome. PLoS computational biology. 2011;7(5):e1001139.

59. Pan RK, Chatterjee N, Sinha S. Mesoscopic organization reveals the constraints governing *Caenorhabditis elegans* nervous system. PloS one. 2010;5(2):e9240.

60. Jarrell TA, Wang Y, Bloniarz AE, Brittin CA, Xu M, Thomson JN, et al. The connectome of a decision-making neural network. Science. 2012;337(6093):437–444.

61. Cook SJ, Jarrell TA, Brittin CA, Wang Y, Bloniarz AE, Yakovlev MA, et al. Whole-animal connectomes of both Caenorhabditis elegans sexes. Nature. 2019;571(7763):63.

62. Yan G, Vértes PE, Towlson EK, Chew YL, Walker DS, Schafer WR, et al. Network control principles predict neuron function in the Caenorhabditis elegans connectome. Nature. 2017;550(7677):519.

63. Csoma A, Kőrösi A, Rétvári G, Heszberger Z, Bíró J, Slíz M, et al. Routes obey hierarchy in complex networks. Scientific reports. 2017;7(1):7243.

64. Avena-Koenigsberger A, Mišić B, Hawkins RX, Griffa A, Hagmann P, Goñi J, et al. Path ensembles and a tradeoff between communication efficiency and resilience in the human connectome. Brain Structure and Function. 2017;222(1):603–618.

65. Akhshabi S, Sarda S, Dovrolis C, Yi S. An explanatory evo-devo model for the developmental hourglass. f1000research. 2014;3.

66. Bargmann CI, Marder E. From the connectome to brain function. Nature methods. 2013;10(6):483.

67. Gray JM, Hill JJ, Bargmann CI. A circuit for navigation in *Caenorhabditis elegans*. Proceedings of the National Academy of Sciences of the United States of America. 2005;102(9):3184–3191.

68. Piggott BJ, Liu J, Feng Z, Wescott SA, Xu XS. The neural circuits and synaptic mechanisms underlying motor initiation in C. elegans. Cell. 2011;147(4):922–933.

69. Tishby N, Pereira FC, Bialek W. The information bottleneck method. arXiv preprint physics/0004057. 2000;.

70. Shwartz-Ziv R, Tishby N. Opening the black box of deep neural networks via information. arXiv preprint arXiv:170300810. 2017;.

71. Lechner M, Grosu R, Hasani RM. Worm-level Control through Search-based Reinforcement Learning. arXiv preprint arXiv:171103467. 2017;.

72. Chalfie M, Sulston JE, White JG, Southgate E, Thomson JN, Brenner S. The neural circuit for touch sensitivity in *Caenorhabditis elegans*. Journal of Neuroscience. 1985;5(4):956–964.

73. Goodman MB. Mechanosensation. WormBook: the online review of C elegans biology. 2006;p. 1–14.

74. Riddle DL, Blumenthal T, Meyer BJ, Priess JR. Mechanosensory Control of Locomotion. In: C. elegans II. Cold Spring Harbor Laboratory Press; 1997.

75. Driscoll M, Kaplan J. Mechanotransduction. In: C. elegans II. Cold Spring Harbor Laboratory Press; 1997.

76. Guo ZV, Hart AC, Ramanathan S. Optical interrogation of neural circuits in *Caenorhabditis elegans*. Nature methods. 2009;6(12):891.

77. Lockery SR. Neuroscience: A social hub for worms. Nature. 2009;458(7242):1124.

78. Li W, Feng Z, Sternberg PW, Xu XS. A *C. elegans* stretch receptor neuron revealed by a mechanosensitive TRP channel homologue. Nature. 2006;440(7084):684.

79. Pirri JK, Alkema MJ. The neuroethology of *C. elegans* escape. Current opinion in neurobiology. 2012;22(2):187–193.

80. Husson SJ, Gottschalk A, Leifer AM. Optogenetic manipulation of neural activity in *C. elegans*: from synapse to circuits and behaviour. Biology of the Cell. 2013;105(6):235–250.

81. Schafer WR. Mechanosensory molecules and circuits in *C. elegans*. Pflügers Archiv-European Journal of Physiology. 2015;467(1):39–48.

